# CD36: Hemin interaction axis to control immune responses and cytokine secretion from macrophages involving Lyn kinase

**DOI:** 10.1101/2021.06.06.447270

**Authors:** Sooram Banesh, Sourav Layek, Vishal Trivedi

**Affiliations:** Malaria Research Group, Department of Biosciences and Bioengineering, Indian Institute of Technology-Guwahati, Guwahati-781039, Assam, India

**Keywords:** Malaria, hemin, biophore, pro-inflammatory cytokines, CD36, signalling

## Abstract

The intensity and duration of TNF-α production are mutually correlated with the level of CD36 expression level. The macrophages exposed to hemin exhibits modulation of non-opsonic phagocytosis of aged RBCs and ability to kill bacteria. Immuno-fluorescence study indicates translocation and sequestration of CD36 within the intracellular storage in the hemin treated macrophages. It in-tern dysregulate the global cytokine secretion from macrophages. CD36 has suitable hemin biophoric environment involving R292, D372 and Q382 to bind and the mutation in biophore residues (R292A, D372A or Q382A) significantly reduced the affinity. Ectopic expression of CD36 in MG63 cells showed several folds increment in cytokines TNF-α, MCP-1, RANTES and CCL1 in response to hemin stimulation but no significant amount of cytokines released with mutants (R292A, D372A or Q382A), highlights the relevance of CD36-hemin interaction for immune-dysfunction. Hemin is driving down-stream signalling involving CD36 and subsequent recruitment of adaptor proteins to the cytosolic domain of CD36. Immuno-precipitation of membrane bound CD36 and detection of adaptor proteins indicate change in level of Lyn proteins with CD36 fractions after hemin stimulation to macrophages. The Lyn targeted siRNA restored the phagocytic activity, reduced the secretion of pro-inflammatory cytokine levels clearly suggests the Src family protein Lyn is crucial for CD36-hemin mediated immune dysregulation and cytokine secretion. In summary, hemin-CD36-Lyn cytokine signalling axis could be a contribution factor to severe malaria pathology and prognosis.

## Introduction

*Plasmodium falciparum*, the causative agent of malaria, is responsible for high mortality rate especially in children and pregnant women (1). During the course of malaria, RBC lysis releases the large amount of pro-oxidant molecules including free hemin and its derivative. During the infection the concentration of hemozoin and hemin immediate after RBC rapture may be as high as 100 µg/ml (2). Macrophages control parasite load through phagocytosis of IRBC in an opsonic or non-opsonic dependent mechanism (3). Phagocytic internalization of IRBC, as well as parasite-derived products, release the large quantity of methemoglobin and hemin inside the macrophages. It is in-turn inhibits the fusion of lysosome to phagosome; disrupts the antigen processing and presentation through partially characterized mechanism.

Macrophage utilizes cell surface and cytosolic receptors to identify infected RBCs (IRBC) and parasite-derived products. Hemin, glycophosphatidylinositol (GPI) and hemozoin activate macrophages to produce cytokines which can excessively fuel inflammatory outcomes (4). Experimental evidences suggest that hemozoin can activate macrophages through TLR9 to produce pro-inflammatory cytokines such as TNF-α. Earlier studies have established that the intensity and duration of TNF-α production are mutually correlated with the level of CD36 expression (5). Moreover, it has been found that macrophage CD36 present on the cell surface binds multivalent ligands, which triggers signal transduction and receptor ligand internalization. Hence it is interesting to explore the effect of hemin on non-opsonic phagocytosis of macrophages, and whether it is related to the CD36 expression on the cell surface and down-stream cytokine secretion.

In this study, we have observed that hemin potentiates drastic changes in the cellular physiology of macrophages such as non-opsonic phagocytosis, ability to kill bacteria and production of pro-inflammatory cytokine TNF-α. Interestingly we have found that hemin exposure not only modulates the pattern of phagocytosis but also leads to sequestration of CD36 receptors within the intracellular storage. The hemin treatment, also induced secretion of series of pro-inflammatory cytokines such as TNF-α, MCP-1, RANTES and CCL1. The pro-inflammatory cytokine secretion upon hemin treatment could explain the pathological outcome during severe malaria. The molecular modelling and affinity studies have confirmed that hemin interacts with the human CD36 ectodomain on a well-defined region consisting of R292, Q382 and D372 residues. The mutations in biophoric residues drastically reduced the affinity and validates the predicted biophore for hemin. We have further, investigated whether TLRs has any role in hemin mediated immune response. The cell line MG63 (with low levels of TLRs and CD36) has been chosen to express CD36 ectopically. The CD36 ectopic expressing MG63 cells migrated towards hemin and confirms the membrane bound CD36 is crucial for hemin to interact and TLRs has no role in it. Besides, the CD36 (wildtype) expressing MG63 cells responded to hemin but mutants (R292A or D372A or Q382A) expressing MG63 cells failed to produce significant levels of cytokine in presence of hemin. Further the hemin treatment activated several proteins including CD36 and suggests the possible downstream signalling. It has been reported that activated CD36 enables the docking of adaptor proteins to cytosolic domain. Our study has found that Src family member adaptor protein Lyn interacts with CD36 in hemin stimulated macrophages as evident from the co-immunoprecipitation study. The Lyn knockdown in macrophages corrected the abnormal phagocytic behaviour and cytokine burst and indicates the Hemin-CD36-Lyn axis is likely to be the reason behind pro-inflammatory cytokine secretion. We have also highlighted that dysfunctional CD36 receptor dynamics (from intracellular vesicles to the cell membrane) may have promising implication in correcting the coherent secretion of cytokines and immunopathology. Our study may provide fundamental understanding role of macrophage in cellular protection, cytokine secretion to contribute into inflammation and brain pathology.

## Experimental Procedures

### Reagents

Peptone Type-III bacteriological, Yeast extract powder, Sodium chloride, ampicillin, skim milk powder, tween-20, dialysis membrane-70, foetal bovine serum were procured from HiMedia labs, India. Isopropyl β-d-1-thiogalactopyranoside (IPTG), Nickel sulfate hexahydrate, Dulbecco’s modified eagle’s medium were purchased from Sigma-Aldrich, India. Site specific primers were procured from Bioserve India, India, and Ni-NTA affinity columns were from Qiagen, Hilden, Germany. Restriction enzymes and 1 kb DNA ladder were purchased from New England Biolabs, USA. PCR master mix (2x) was purchased from Fermantas, India. Dual color proteins standards purchased from Biorad, California, USA. T4 DNA ligase was purchased from genetix, India. Desalting column (PD-10) was purchased from GE life sciences, India. Human CD36 gene, mApple-CD36-C-10 was a gift from Michael Davidson (Addgene plasmid # 54874). The pET23a vector, BL21-(DE3), DH5α bacterial strains were from novagen. Mouse polyclonal anti-6xHis antibody, rabbit polyclonal anti-phosphotyrosine antibody purchased from Sigma-Aldrich, India. Rabbit anti-CD36 antibody, Mouse anti-CD36 antibodies rabbit HRP conjugated secondary antibody were procured from Santa Cruz Biotechnology, USA. ECL reagent was procured from Bio-Rad labs, USA. DMEM-F12, DMEM-high glucose are from Sigma-Aldrich. PEiRfect transfection reagent was procured from Biobharati lifesciences, India. The cytokine ELISA kits for TNF-α (cat#. KRBA10602-5), MCP1 (cat#.KRB10134-5) and RANTES (cat#.KB1102) were purchased from Krishgen Biosystems, Mumbai, India and CCL1 (cat#.KTL11769) from (Kreative Technolabs, Los Angeles, CA, USA)

### Mammalian cell culture

Different cell lines J774A.1, MG63, MDAMB-231, A549, HeLa, HEK-293, SAS, Sf-9, and Sf-21 were purchased from NCCS, Pune, India. All the cell lines were maintained in DMEM high glucose medium supplemented with 10% of Foetal bovine serum containing 1% penicillin and streptomycin at 37°C in a humidified chamber with 5% CO_2_.

### Phagocytosis assay

Macrophages were either remain untreated or treated with different non-toxic concentration of hemin (0-200µM) in serum free medium for 1hr at 37°C in a humidified chamber with 5% CO_2_. Post treatment, hemin was washed three times with PBS and cells were removed from the culture dish by incubating with 0.5% EDTA for 30 mins and phagocytosis was measured in flow cytometry-based assay as described (6). In the assay, cell suspension was incubated with FITC-labelled E. Coli (1:10) for 1hr at 37°C with intermittent shaking and then placed on ice. Trypan blue (1%) was mixed with the cells to quench the fluorescence from the bacteria sticking to the cells. Ten thousand cells were analysed by flow cytometry (FL1 channel) using FACS Calibur (Becton-Dickson) to calculate the proportion of cells phagocytose bacteria whereas mean fluorescence intensity (MFI) was used to calculate number of bacteria taken up per cell. Data is expressed as phagocytic index using the formula; phagocytic index= % phagocytic macrophages X MFI.

To study the phagocytosis of different objects in the microscopy based assay, normal RBC (NRBC), oxidized RBCs (oxi-RBC), latex beads coated with LPS, Amine beads (+Ve charged) were incubated with untreated or hemin (75µM) treated macrophages for 1hr at 37 °C. Cells were washed with PBS to remove un-phagocytosed objects and stained with fluorescent dye filipin (5µg/ml) to identify the phagosomes. Cells were observed under the fluorescence microscope 80i (Nikon) and random 10 different fields were selected to count number of phagosomes to calculate the phagocytic activity of macrophages.

### Bactericidal Assay

Bactericidal assay was performed as described previously (7). Hemin treated macrophages (10^6^) were incubated with live *E.coli* suspension (1:50) for 1hr at 37 °C. After 1hr, tubes containing the cell suspension were centrifuged (300×g) for 12mins at 4°C. Cell pellet was washed thrice with PBS to remove unbound bacteria and macrophages were lysed with chilled milli-Q water. Cell lysate was added to the 3ml LB and allowed to grow at 37°C overnight, number of bacteria were counted and used to calculate bactericidal activity.

### Immunolocalization of CD36 in macrophages

The murine macrophages J774A.1 were grown on a cover slip (in a 24 well plate) and treated with hemin (0, 25 or 200 µM) for 1 h in serum free medium. The cells were washed with PBS to remove free hemin then fixed and permeabilized. The cells were incubated with blocking solution (3% BSA in PBS) for 1 h in CO_2_ incubator followed by anti-CD36 antibody (1:500 dilution) incubation at 4 °C for 4 h. The cells were washed with PBS to remove any unbound antibody and incubated with FITC conjugated secondary antibody (1:700 dilution) for 2 h at 4 °C. The secondary antibody removed and washed with PBS. The coverslip was mounted on a glass slide and observed under Nikon fluorescence microscope. The images for bright field or fluorescence were collected from ten random fields and analysed.

### Macrophage cell fractionation and assessment of purity

The macrophages seeded at 5 x 10^6^ per well in a 6 well plate. The cells were either remain untreated or treated with hemin (25 or 200 μM for 1 hr at 37 °C. Subsequently, macrophages were fractionated to prepare cytosol and membrane as described previously (8). The membrane and cytosolic fractions were assessed for their purity by their respective marker enzyme assay.

### Cytokine profiling

Macrophages J774A.1 were either remain untreated or treated with hemin (25µM) in serum free DMEM medium for 1hr and subsequently cells were allowed to secrete cytokines overnight (∼18 hrs) in cell culture supernatant at 37 °C in humidified CO_2_ incubator. The cytokines released from macrophages treated with hemin were detected by mouse cytokine proteome profiler array panel A kit (ARY006, R&D Systems, USA) as per the manufacturer’s instructions. The dots corresponding to different cytokine signals were analysed using densitometry (ImageJ, National Institute of Health, USA) and normalized to control dots present in the array.

### Measurement of cytokines in cell culture supernatants

The murine macrophages maintained in DMEM medium were seeded at 2 x 10^4^ cells per well in a 96 well plate and allowed to adhere. The cells were washed and treated with hemin (prepared in a serum free medium) for 1 hr. Post-treatment, cells were washed with PBS and the serum free medium was added to the cells and allowed to secrete the cytokine overnight at 37^0^C in humidified CO_2_ incubator. The cell culture supernatants were collected and centrifuged to remove the debris and dead cells. TNF-α level was measured using ELISA kit (555268, BD Biosciences, Sa Jose, CA, USA). In experiments with human osteosarcoma (MG63) cells transfected with CD36_cherry, different cytokines such as TNF-α (KRBA10602-5, Krishgen Biosystems, Mumbai, India), MCP1 (KRB10134-5, Krishgen Biosystems, Mumbai, India), RANTES (KB1102, Krishgen Biosystems, Mumbai, India), and CCL1 (KTL11769, Kreative Technolabs, Los Angeles, CA, USA) level in the cell culture supernatant were quantified as per the manufacturer’s instructions. All the data was corrected for substrate absorbance and plotted using Origin pro software (Origin labs, Northampton, MA, USA). The levels of cytokines were expressed in ‘folds’ change considering the level of cytokine present in ‘untreated’ cells.

### Prediction of Hemin biophore in CD36

The hemin co-crystalized proteins (1O9X, 2BLI, 2QSP, 2R7A, 2ZDO, 3NU1, 4MF9 and 4MYP) were downloaded from protein data bank (www.rcsb.org) and analyzed for binding environment of hemin within 6 Å using protein contact map webserver ContPro (9). The interactions between protein and hemin atoms were analysed using Ligplot+ and protein-ligand interaction profiler webserver (https://projects.biotec.tu-dresden.de/plip-web/plip/). The residues found consistently featuring in majority of the analysed complexes were mapped and a matrix has been generated with hemin and protein atoms. The distance between atoms scaled and colour coded in VIBGYOR.

### Generation of CD36-Hemin molecular model

The Autodock v4.2 with autodock tools v1.5.6 application with genetic algorithm as a scoring function was used for CD36-hemin molecular docking (10). The molecular docking was performed on predicted biophore as grid centre. A total of 100 GA runs with 2500000 energy evaluations were carried out. The docking log file was analysed for binding energy, number of clusters and conformations in each cluster. The best pose with highest conformations and lowest binding energy was analysed for hemin-CD36 interactions using ligplot+ and protein-ligand interaction profiler webserver application.

### Molecular dynamics simulations

The crystal structure of CD36 (PDB id: 5LGD) was downloaded from protein data bank and mutants were generated using modeller. The topology file for hemin was generated based on the topographical inputs from the webservers PRODRG (11) and automated topology builder (ATB) (12). The MD simulations were performed on Dell precision T1700 workstation using GROMACS 4.6.5 package. The GROMOS96 43a1 force field was used for simulations. The protein-hemin complex was placed in the simulation box such that the complex is 1.5 nm distant from the wall of the simulation box. The system was solvated with simple point charge water and energy minimized by steepest descent algorithm. The production run was performed for 10 ns for wild-type, and mutational variants (R292A, D372A and Q382A) with an integration step of 1 fs under NVT conditions. LINCS algorithm with geometric accuracy of 10^−4^ was used as the bond length constraint. Maxwell distribution was used for initial velocity calculations with 0.1 ps of coupling relaxation time at 300 K. The non-bonded interaction cut off was set at 1.4 nm for both van der Waal’s and electrostatics (PME). The MD trajectories after production run were analysed for root mean square deviation (RMSD), structure-based clustering, and radius of gyration (Rg), root mean square fluctuation (RMSF).

### Generation of Membrane Bound CD36

For membrane bound CD36 molecular dynamics study, a 128 membered Dipalmitoylphosphatidylcholine (DPPC) lipid bilayer was constructed using the MemBuilder server (13). GROMOS 53a6 force field parameters were used in combination with Berger lipids force field. Water as simple point charge was used as the solvent. Parrinello-Rahman barostat was used was used for pressure coupling and isothermal compressibility was set at 4.5 × 10^−5^ bar^−1^. The system was energy minimized by the steepest descent algorithm and equilibrated for one nano second (ns) under NVT (conserved <u>N</u>umber of particles, <u>V</u>olume and <u>T</u>emperature) and NPT (conserved <u>N</u>umber of particles, <u>P</u>ressure and <u>T</u>emperature) conditions successively. A total of 100 ns production run was performed on the system containing CD36-hemin integrated into DPPC bilayer. Post-simulation, the system was analysed for the bilayer thickness, RMSD (root mean square deviation) and Rg (radius of gyration).

### Cloning, over-expression and purification of hCD36ecto wild type and mutants

The cloning, overexpression, and purification of hCD36ecto was done as reported in our previous publication (14). Different mutants were also prepared following the similar procedure.

### Site-directed mutagenesis

The conventional long-range PCR amplification (15) performed to introduce mutations at R292 (R292A), D372 (D372A) and Q382 (Q382A) positions in pET23a-hCD36ecto (for expression in bacterial system) or pmApple-CD36-C-10 (for expression into mammalian system). The different primers used for generation of mutant CD36 is given in Table S1. The PCR reaction was performed in a total volume of 50 μL containing 20 ng of template (pET23a-hCD36ecto or pmApple-CD36-C-10), 15 pM of mutagenic primers (Table S1) and PCR master mix. The PCR amplification was carried out at 94 °C initial denaturation for 4 min followed by 16 cycles of denaturation (at 94 °C for 30 s), annealing (60 °C for 1 min) and extension (68 °C for 7 min). The amplified products were PCR cleaned up and eluted in nuclease-free water. Subsequently, the eluted product digested with *Dpn-I* for 2 h at 37 °C. Finally, the digested product transformed into NEB 5α competent cells (New England Biolabs, Ipswich-MA, USA). The mutation was confirmed by sequencing and restriction digestion.

**Table 1:** Effect of hemin on Macrophage for phagocytosis of different ligands.

| Phagocytic Index |  |  |  |  |
| --- | --- | --- | --- | --- |
| S.No. | Ligand | Untreated ±SD | hemin treated ±SD | $\Delta$ Phagocytic index ±SD |
| 1 | +Ve Charged | 87.27 ± 3.26 | 39.53 ± 3.289 | 45.29 ± 3.277 |
| 2 | LPS | 86.71 ± 9.08 | 43.57 ± 10.833 | 49.76 ± 9.958 |
| 3 | NRBC | 73.29 ± 13.61 | 40.55 ± 5.250 | 44.67 ± 3.417 |
| 4 | PS containing RBC | 71.39 ± 9.09 | 25.20 ± 2.602 | 64.70 ± 4.876 |

Foot note: Macrophage were treated with hemin (75 μM) for 1 hr at 37<sup>0</sup>C in serum free media. Hemin was washed with PBS and allowed to phagocytose normal RBC (NRBC), PS containing RBC, amine beads (+ve Charged) and LPS coated beads (LPS) for 1hrs at 37<sup>0</sup>C in serum free media and phagocytosis was measured and expressed as phagocytotic index ± SD. (n=4)

### Dot blot assay

The dot-blot assay was performed as described (14). Briefly, the hemin solution (50µM) was applied to a nitrocellulose membrane blot as a dot and air-dried. The membrane was incubated with the blocking buffer containing 3% BSA in TBST buffer (50 mM Tris-HCl, 150 mM NaCl, 0.05% Tween-20, pH 7.5) for 2 hr at room temperature. The membrane was incubated with purified hCD36ecto for 2 hr at room temperature with mild shaking on a shaker. Subsequently, the NC membrane was washed with TBST buffer to remove any unbound proteins and incubated with anti-CD36 antibodies for 2 h at room temperature. After washing, the blot was incubated with HRP-conjugated secondary antibody to detect the presence of CD36 on the membrane. Dot blot was developed using Clarity western ECL substrate (1705061, Biorad labs, California, USA) and images were acquired in the ChemiDoc MP imaging system (Biorad labs, USA). Similarly, the assay was performed on mutants (R292A, D372A, and Q382A) as well to test the affinity of hemin to the mutant proteins. The signal intensity was normalized or calculated by considering wild type as 100 %.

### Isothermal titration calorimetry

The affinity measurements of hemin towards wild type (hCD36ecto) or mutants (R292A, D372A and Q382A) were performed in MicroCal iTC200 isothermal titration calorimeter (GE Healthcare, Chicago, USA). Unless specified, all the titrations were carried out at 37°C. The wild type or mutant proteins (0.01 mM) were loaded into ITC titration cell in a total volume of 200 μL. The syringe is filled with hemin solution (0.1 mM) and placed in ITC titration cell. A total of 25 injections were carried out consisting of 0.4 μL of first injection and 1.5 μL in subsequent injections. To correct background heat generated due to buffer or hemin dilution, another blank titration was carried out with hemin against buffer only. The data were analyzed using ITC200 analysis software v7.2 application integrated in ORIGIN7 graphing software (Origin Lab, USA) and fitted with sequential binding mode to calculate the dissociation constant.

### Transfection studies

All the transfections were performed according to the manufacturer’s instructions. The MG63 cells with low passage number were seeded at 3 x 10^5^ cells per well in a 6 well plate one day prior to the transfection. The wild type (pmApple-CD36-C-10) or mutant (pmApple-R292A, pmApple-D372A and pmApple-Q382A) plasmids were diluted in 50 μL of serum free DMEM medium and vortexed briefly for 5 sec. The PEI transfection reagent was added to diluted plasmid DNA at 1:2 ratio (2 μg of PEI for 1 μg of plasmid). The DNA+PEI mixture vortexed for 5 sec and incubated at 25^0^C for 15 min to form the DNA-PEI complexes. The transfection mixture was added drop-wise to the cells and after 6 hr, medium was replaced with DMEM complete medium and incubated for another 48 hr for expression. The expression was analysed by fluorescence imaging, flow cytometry [using (FL-2 channel, λex488/λem575)] and western blotting.

### Chemotaxis assay

The chemotaxis assay was performed as described (16). The transfected MG63 cells seeded at 2 x 10^4^ cells in serum free DMEM in upper chamber of inserts (with 12 μm pore size) in a 24 well dish. The bottom chamber is filled with either serum free medium or hemin (prepared in serum free medium). The cells were kept in a CO_2_ incubator at 37 °C for 6 h to allow the migration. After 6 h incubation, the inserts were washed with PBS to remove dead cells and floating debris and fixed in 95% ice-cold ethanol for 8 min. The inserts were stained with hematoxylin and eosin successively. The inserts were washed with the PBS to remove the excess dye and the cells in upper chamber side were removed by gently scraping with cotton swab. The cells in lower chamber side were observed and photographed in ten random fields. The chemotactic index was calculated from the formula given below:

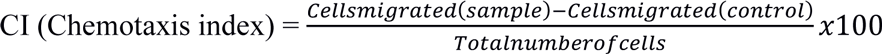

### Immuno-precipitation

The J774A.1 cells were seeded at 5 x 10^7^ in a 100 mm cell culture dish one day before the experiment. The cells were treated with hemin (25 µM) for 1 h in serum free medium and washed three times with PBS. The cells were detached using cell scraper and the cell pellets was re-suspended in IP lysis buffer (10 mM HEPES pH 7.4, 250 mM NaCl, 1 mM EDTA, 0.5% NP-40 and 0.2% Tween-20) supplemented with freshly prepared protease (1 mM PMSF) or phosphatase inhibitors (10 mM NaF and 0.1 mM Na_3_VO_4_). The cell suspension passed through 2-bend 25-gauge needle ten times to homogenize the lysate and incubated on ice for 20 min. Subsequently, the lysates were clarified by centrifuging at 14,000 rpm for 2 min at 4°C. The protein concentration was estimated using Bradford assay (17). The clarified lysates was pre-cleared by incubating with 20 µL of protein AG plus agarose beads slurry (BB-PAG001PA, Biobharati lifesciences, Kolkata, India) at 4°C for 1 hr under rotating condition. The pre-cleared lysates were incubated with mouse anti-CD36 antibody (2 µg) or anti-Lyn antibody (1:50 dilution) at 4 °C overnight under rotating condition. The pre-washed 20 µL of protein AG agarose beads were added to the lysate-antibody mixture and incubated overnight at 4°C on a rotator. The protein AG agarose beads were washed with IP lysis buffer four times and eluted in SDS-PAGE sample buffer containing 150 mM of DTT.

### Analysis of the phospho signal in hemin treated macrophage cell lysates

The J774A.1 cells maintained in DMEM high glucose with serum supplementation were seeded in a 100 mm cell culture dish one day before the experiment. The cells were incubated with indicator free DMEM and kept for 4 h in incubator to eliminate the phenol red induced phosphorylation in cells. After 4 hr, the cells were treated with hemin (25µM or 200 µM) in indicator free DMEM for 10 min. The cells were lysed in 10 mM HEPES, pH 7.4 buffer and the protein concentration was estimated using Bradford assay. The 50 µg lysate was resolved in 10% SDS-PAGE and transferred onto polyvinylidene difluoride (PVDF) membrane using wet transfer method. The blots were incubated with blocking solution (5% BSA in TBST buffer, pH 7.4). Post-blocking, the membranes were incubated with anti-phosphotyrosine followed by HRP conjugated secondary antibody. After washing with TBST buffer, the blots were developed using chemiluminscence substrate. The blots were analysed for molecular weight and number of phosphor-reactive protein bands using Biorad’s Imagelab software V.6.0.1. The molecular weight of the bands was compared with the phosphosite-plus server (https://www.phosphosite.org/homeAction.action) for probable proteins involved in signalling event.

### Statistical analysis

All experiments unless otherwise stated were performed in triplicate and repeated at least three times. All the data are presented as the mean ±SD unless otherwise noted. Differences between groups were analyzed using the sigmaplot (Systat software inc, San jose, CA, USA) and Originpro (Originlabs, Northampton, MA, USA) data analysis and graphing software. Statistical analysis was performed by one-way ANOVA followed by unpaired Student t-test and the p < 0.05 were considered as statistically significant.

## Results

### Hemin exposure Disturbs macrophage phagocytosis towards oxidized RBCs

Hemin is a toxic pro-oxidant molecule but at higher concentration and longer exposure time periods (18). Hemin exposure to macrophages at different concentrations (0-300µM) in serum free or complete medium for 1hr at 37^0^C in humidified CO_2_ incubator is not causing reduction in cellular viability (Figure S1). Almost more than 90% viability was observed for macrophages treated with hemin concentration up- to 200µM and this range is been used for experiments throughout current work. Macrophages treated with different concentrations of hemin (0-200µM) exhibits phagocytic activity with a unique pattern in a flow-cytometry based phagocytosis assay (Figure 1A and Figure S2). At lower concentration (25µM), hemin stimulates macrophage phagocytosis, followed by dose-dependent depression of phagocytic activity at higher concentrations (75 µM). A similar pattern of enhancement followed by depression of phagocytic activity of hemin treated macrophages continued with complete loss of phagocytosis at the highest concentration (200 µM). Interestingly, the ability of hemin to stimulate phagocytic activity was reduced with increasing hemin concentration resulting in loss of phagocytosis at higher hemin concentration (Figure 1A). The phagocytosis pattern indicates the macrophages with different phagocytic activity; (1) Macrophage treated with low hemin concentration (25 µM) exhibiting high phagocytic activity (2) macrophage treated with high hemin concentration (75 µM) with very low phagocytic activity (Figure 1A). Macrophages phagocytose target objects with different types of ligands either in non-opsonic or opsonic manner (19). We further tested the phagocytic activity of hemin treated macrophages towards different ligands. Hemin exposure caused global depression of phagocytic activity of macrophages towards amine, LPS and phosphatidylserine containing oxidized RBCs (Table 1). After careful observation of these cells under the fluorescence microscope, it was established that the depression in macrophage phagocytosis is more severe towards oxidized RBCs (containing PS) compared to the normal RBCs (Figure 1B and Table 1). Phagocytosis is followed by internalization of the infectious agent and its destruction within phagolysosomes through the action of lytic enzymes (20). Hemin treated macrophages were showing severe reduction in bactericidal activity compared to the untreated cells (Figure 1C).

**Figure 1:**
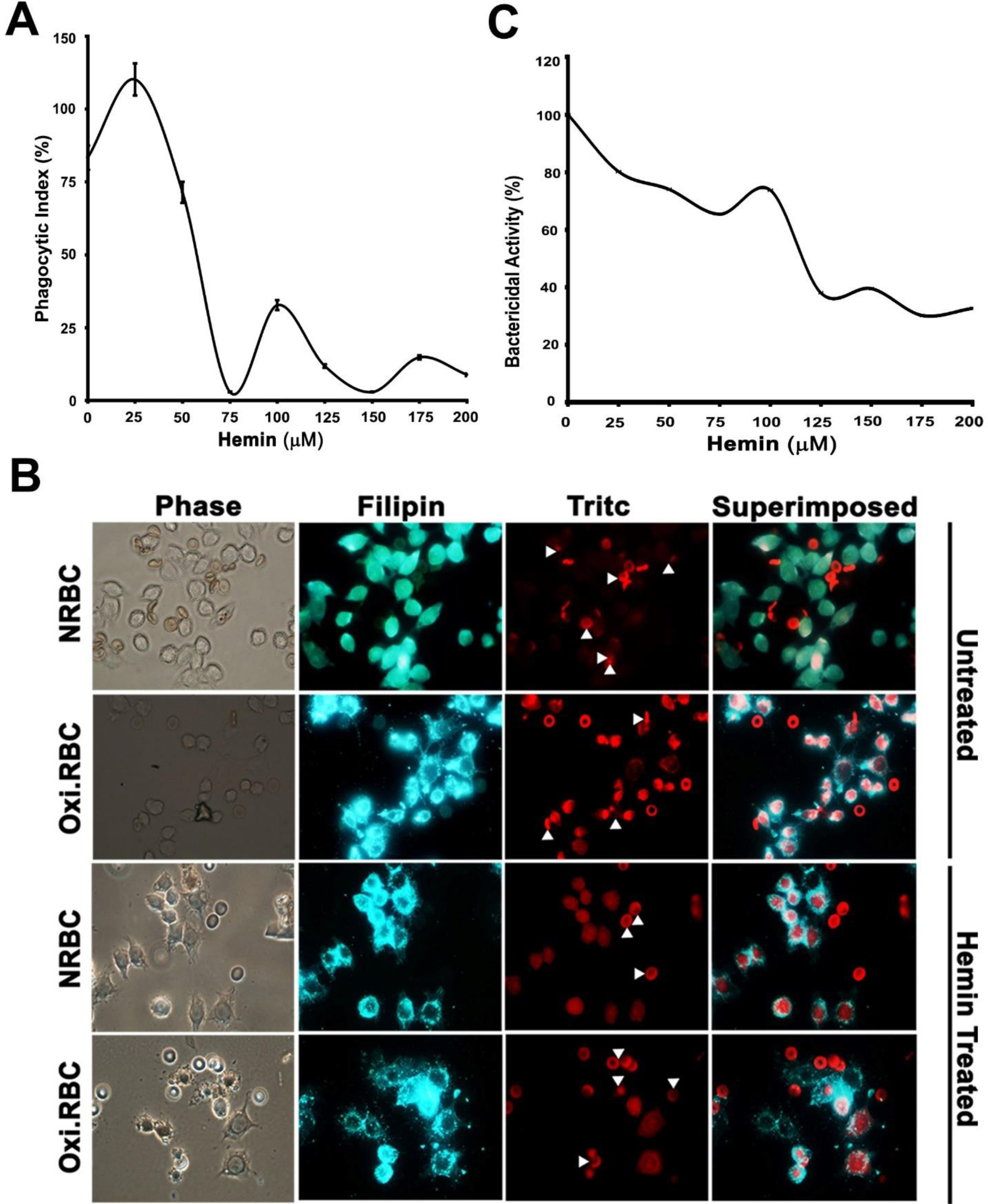
Macrophages treated with hemin exhibits phagocytotic defects. **(A)** Macrophages treated with different concentration of hemin (0-200µM) and phagocytotic activity was measured in flow-cytometry based assay as described in “material and methods”. n=3 **(B)** Depression in phagocytosis of hemin treated macrophage is more severe towards oxidized RBCs (containing PS) compared to the normal RBCs. Macrophage were treated with hemin (75 µM) for 1 hrs at 37 ^0^C in serum free media. Hemin was washed with PBS and allowed to phagocytose normal RBC (NRBC), oxidized RBC (Oxi.RBC). Macrophages were stained with filipin (Blue colored fluorescence dye to identify phagosomes containing engulfed material) and observed using the fluorescence microscope Nikon 80i in DAPI and TRITC filters to acquire filipin (Blue) and RBC (Red) fluorescence respectively. n=3 (C) The bactericidal activity of macrophages treated with hemin was drastically reduced. At low hemin (25 µM) the bactericidal activity was found to be 80% compared to untreated whereas the actvity reduced with increase in hemin concentration.

### Hemin directs the sequestration of membrane bound CD36 into intracellular vesicles

We have two macrophage populations; macrophage with high phagocytosis activity at 25µM (subset 1) and low phagocytosis activity at 200µM (subset 2). CD36 is the key receptor responsible for non-opsonic phagocytosis of PS containing dead and aged RBCs (21). Presence of CD36 on macrophage plasma membrane is crucial for phagocytosis of RBCs to facilitate the clearance from circulation. CD36 level on the plasma membrane was measured on both macrophage subsets to explore the role of the receptor with the observed change in phagocytosis activity of macrophages. Subset-1 macrophages (Cells treated with low hemin, 25µM) exhibit increased level of CD36 on plasma membrane whereas low level of CD36 was found on subset-2 (Cells treated with high hemin, 200 µM) cells (Figure 2A, inset). CD36 localization inside the cells indicates the high level of CD36 on the vesicular structures especially in the subset-2 cells and major sequestration of CD36 within the vesicular structures. It may probably be responsible for CD36 down-regulation on the plasma membrane in these cells (Figure 2B, inset). Internalization and uptake of hemin were observed in the macrophages treated with different concentration of hemin (data not shown). Interestingly, a large quantity of hemin was observed within the intracellular vesicles. The CD36 sequestration was further confirmed by western blot (Figure 2C). The membrane and cytosol fraction prepared from macrophage is pure and doesn’t have any croos contamination (Figure S3). CD36 signal from the untreated membrane fraction is very intense comparing to the cytosol whereas in the hemin treated macrophages the cytosolic CD36 signal is intensified. The densitometry analysis indicate ∼44.6% of the CD36 was translocated into cytosol. The western blotting data confirms the CD36 sequestration into cytosol.

**Figure 2:**
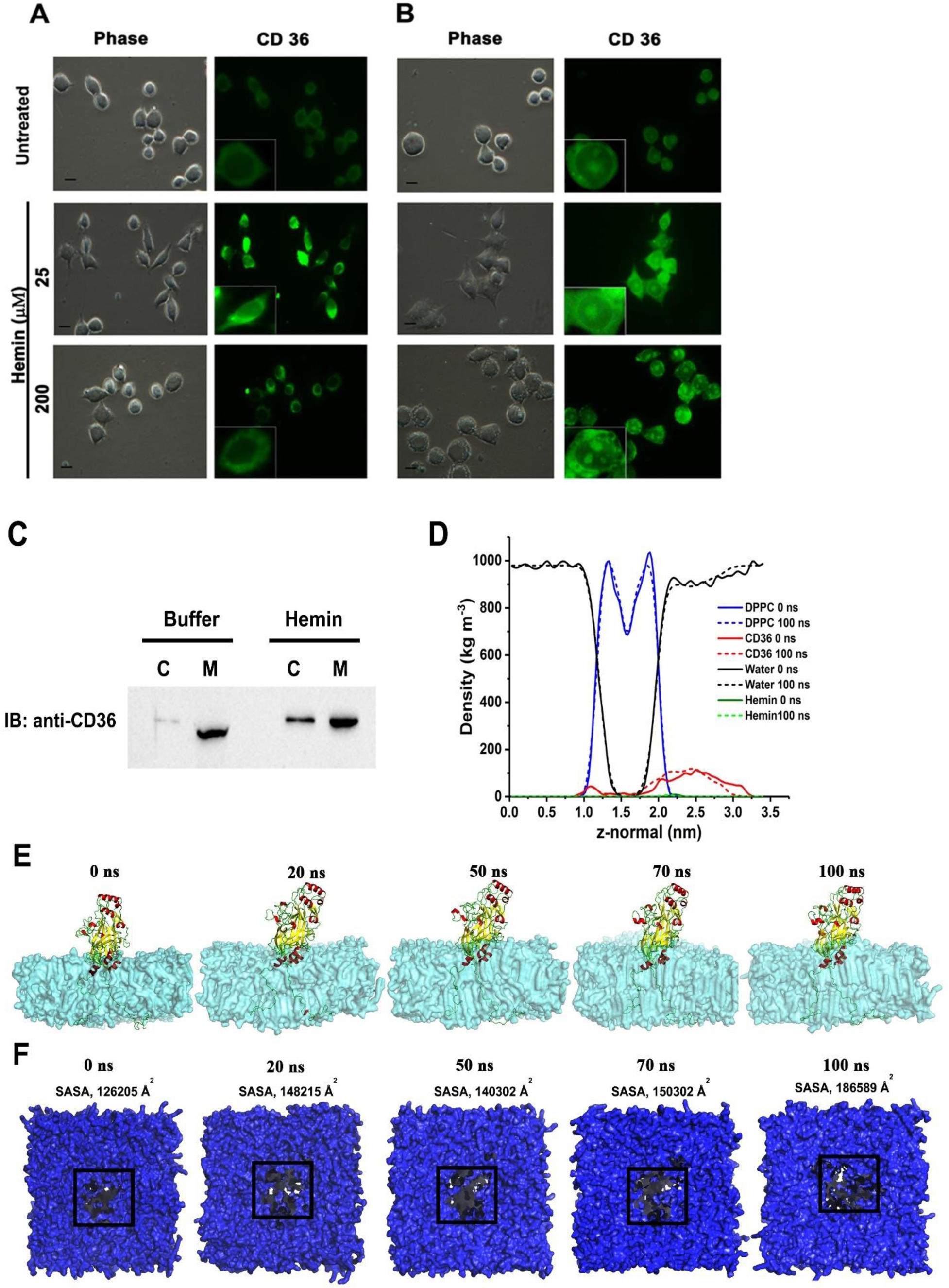
CD36 sequestration into intracellular vesicles in macrophages is linked to hemin interaction with scavenger receptor CD36. Macrophages were stimulated with hemin (25 or 200µM) and stained with anti-CD36 in **(A)** non-permeabilized and **(B)** permeabilized cells to immuno-localize CD36 on the cell surface and intracellular vesicles; (n=4). In permeabilized hemin treated cells, the CD36 signal is localized to cytosolic vesicles. **(C)** Macrophages were treated with hemin and fractionated into membrane and cytosolic fractions and immunoblotted with anti-CD36 antibody. In hemin treated cytosolic fractions, the CD36 signal intensified as compared to the cytosolic fraction from untreated macrophages. **(D)** Hemin binding facilitates the CD36 passage through DPPC bilayer. The MD run trajectories of membrane bound CD36-hemin complex was analysed for density distribution DPPC, CD36, Hemin and water at the starting (0 ns) and end point (100 ns) of the simulation. The CD36 integrated into membrane was penetrated into DPPC bilayer. **(E)** The time lapse membrane bound CD36-hemin complex. **(F)** The top view of DPPC in a time lapse scale. The solvent accessible surface area (SASA) was measured at 0 ns, 20 ns, 50 ns, 70 ns and 100 ns. The thinning of membrane was observed over time frame as evident from SASA. The overall study indicates the hemin binding may facilitate the CD36 passage through DPPC bilayer and supports the CD36 translocation in hemin treated macrophages.

To understand the process, we have developed the membrane bound CD36 receptor bound to hemin and performed molecular dynamic simulation as described in experimental procedure. After successfully integrating the complex into membrane, the system was successively equilibrated in NVT and NPT conditions. A production run for 100 ns was carried out. The structural distortions in CD36-hemin complex during the simulation run was analysed using root mean square deviation (RMSD) and radius of gyration (Rg). The DPPC and water densities were found less affected through the simulation run whereas CD36 bound to hemin showed penetration into the DPPC bilayer (Figure 2D). To assess further, the time lapse frames of simulation run were captured at 0, 20, 50, 70 and 100 ns. The time lapse images suggests the lower region of ectodomain was found to be penetrated in upper leaflet of DPPC (Figure 2E). Further the solvent accessible surface area (SASA) was calculated on bilayer without protein-ligand complex. A significant area (186589 Å^2^) was found to be accessible by solvent at 100 ns compared to initial structure (Figure 2F). Overall, results indicate the thinning of the bilayer over the simulation and indirectly support the penetration of CD36 into the membrane in hemin bound form.

### CD36-Hemin interactions likely to be a contribution factor for pathological outcome during severe malaria

Receptor recycling from intracellular vesicles to the plasma membrane is vital for proper stimulation and regulation of down-stream events such as cytokine secretion (22). The CD36-hemin complex sequestration into intracellular vesicles may contribute to the pathological outcome. Besides, during severe malaria, the free heme levels are very high, and the previous studies suggest the heme levels are correlating with the severity of malaria, organ damage, and the cytokine levels (23). To assess the impact of CD36 sequestration on the secretion of different types of pro-inflammatory or anti-inflammatory cytokines, we have treated macrophages with hemin (25 µM) or 200 µM and the secreted cytokines were detected using mouse cytokine array profiler kit as described in experimental procedure. In untreated macrophage cell culture supernatants, the cytokines such as IP-10, KC, CCL3, CCL4, CXCL2 found to be constitutively expressing whereas very low reactivity was observed for cytokine/chemokines such as G-CSF, GM-CSF, CCL-1, IL-7, IL-16, CXCL-1, MCP-1, TIMP-1 and TNF-α (Figure 3A (top panel), and Figure 3B). The presence of pro-inflammatory cytokines in untreated could be due to the cell debris in cultures which could activate the macrophages to produce cytokines (24). Further, the cytokines/chemokines such as CS/C5a, sICAM-1, IFN-γ, IL-1α, IL-1β, IL-1ra, IL-2, IL-3, IL-4, IL-13, IL-17, IL-23, IL-27, M-CSF, CXCL9 and TREM-1 were found exclusively in hemin treated macrophage cell culture supernatants but not in untreated (Figure 3A, 25 µM panel). In 25 µM hemin treated macrophage cell culture supernatants the up-regulation of pro-inflammatory cytokines such as TNF-α (32 folds), RANTES (7 folds), MCP-1 (2.4 folds), IL-16 (7.5 folds), IP-10 (3.4 folds), GM-CSF (10.6 folds), CCL-1 (3 folds), IFN-γ (1 fold) CS/C5a (1 fold) was observed (Figure 3B). The CS/C5a is a complement protein involved in recruitment and activation of polymorphonuclear cells to site of infection and contribute into inflammation. In experimental cerebral malaria model, the knockdown of CS/C5a showed improved survival of mice. The knockdown is associated with reduced levels of MCP-1, IFN-γ, and TNF-α levels (25). Interestingly, the CS/C5a is present in hemin treated macrophage cell culture supernatants but not in untreated. It has been shown that the cytokines IFN-γ, TNF-α, and IL-12 are crucial for development of CM (26). The knock down of IFN-γ in mice showed no signs of cerebral malaria compared to wild type suggests IFN-γ plays crucial role in development of cerebral malaria pathology (27). The absence of IFN-γ in untreated macrophage cell culture supernatants but in hemin treated macrophages imparts hemin could be a potential pro-oxidant molecule in development of cerebral malaria pathology. Surprisingly, few cytokines IL-17 (5.3 folds), IL-23 (13.2) and M-CSF (3.2 folds) are up-regulated in 200 µM hemin treated macrophages but not in 25 µM (Figure 3B). The cerebral malaria patients with multiple organ damage especially acute renal failure (ARF) distinctly showed elevated levels of cytokines IL-17, IL-10 and IP-10 (28). The study underlined that IL-17 is crucial for development of ARF pathology associated with malaria but the mechanistic details yet to ascertained (28). The cytokine array results clearly indicate that the multiple organ damage and the pathology associated with cerebral malaria could be due to hemin interaction with various receptors present on macrophages or endothelial cells on brain. The pro-inflammatory cytokine TNF-α plays a crucial role in the pathophysiology of the host during cerebral malaria (29). So we further examined the secreted TNF-α levels in response to hemin treatment using ELISA. Macrophages treated with different concentrations of hemin (0-200 µM) show dose-dependent secretion with the highest TNF-α level at 75 µM (phagocytic activity at this concentration is lowest). Subsequently, TNF-α levels are reducing again, rising dose-dependently up to high hemin concentration 200 µM (Figure 3C). Moreover, macrophages stimulated with bacteria were giving several folds high TNF-α secretion compared to the unstimulated cells. Whereas, cells treated with hemin and stimulated with bacteria did not show much TNF-α secretion (Figure 3D). It indicates a functional defect in the hemin treated macrophages towards immunological stimuli to secrete TNF-α for clearing the infection from the site of action. In addition, TNF-α secretion kinetics indicates that the untreated cells stimulated with bacteria were giving rapid TNF-α release (which reduces back to basal level in the next 4 hrs) compared to the hemin treated macrophages under identical conditions (Figure 3E).

**Figure 3:**
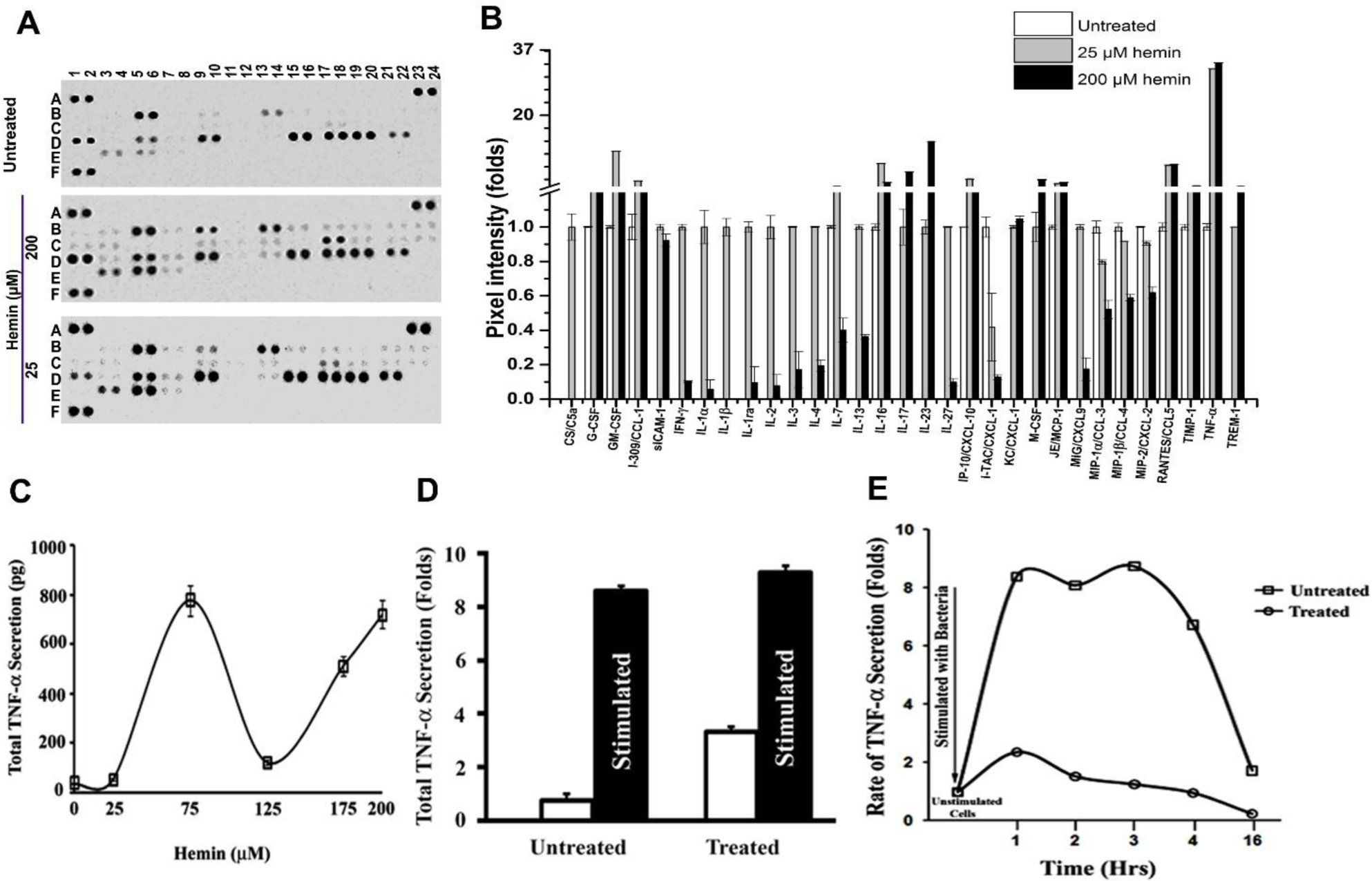
Hemin treatment upregulates various pro-inflammatory cytokines in macrophages. Macrophages were stimulated with various concentrations of hemin (in µM) and analysed for cytokine secretion. **(A)** The macrophages either treated with hemin (25 or 200 µM) or remain untreated were allowed to secrete the cytokines overnight and the next day the cell culture supernatant were collected. The culture supernatants were assessed for the presence of cytokines using muse cytokine array panel A kit. The upper blot (untreated) shows the presence of various constitutively expressing cytokines along with control spots and a negative control spot. The hemin treated cytokine arrays (middle (25 µM) and bottom blot (200 µM) showing upregulated cytokines such as CCL1, IL-16, CXCL1, RANTES, MCP-1 and TNF-α. **(B)** The cytokine dot intensity was semi quantified and normalized by control spots provided in the array. The cytokine levels plotted as normalized pixel density (folds change) calculated using ImageJ application (NIH, USA). The cytokine such as TNF-α, RANTES and MCP1, G-CSF, CCL1 and sICAM1 showing upregulated. **(C)** Macrophages were treated with different concentration of hemin (0-200µM) for 1hrs at 37^0^C in serum free media. Cells were washed with PBS and allowed to secrete TNF-α for 12hrs in serum free media. TNF-α secretion from the cells were measured using ELISA as per manufacturer’s instruction and expressed as pg ±SD. n=3 **(D)** Macrophages either remained untreated or treated with hemin (75 µM) for 1 hrs at 37^0^C in serum free media. Cells were washed with PBS and stimulated with LPS (1µg/ml) for 1hrs, washed to remove LPS and allowed to secrete TNF-α for 12hrs in serum free media. TNF-α secretion from the cells was measured using ELISA as per manufacturer’s instruction and expressed as fold change. **(E)** Macrophages either remained untreated or treated with hemin (75 µM) for 1hrs in serum free media. Cells were washed with PBS and stimulated with LPS (1 µg/ml) for 1hrs, washed to remove LPS and allowed to secrete TNF-α for different time period (1-16 hrs) in serum free media. At each time point, culture supernatant was collected and TNF-α level was measured using ELISA as per manufacturer’s instruction and expressed as fold change.

### Hemin serves as ligand for CD36

The scavenger receptor CD36 interactions with its ligands often induce the internalization of CD36 (30). It has been shown that the binding of α-Tocopherol to CD36 induces CD36 internalization. Further, the use of selective inhibitor of lipid transport by CD36 prevented the α-Tocopherol mediated CD36 internalization (31). The sequestration of CD36 into vesicles and defect in phagocytosis upon hemin treatment lead us to hypothesize that the hemin could be a ligand for CD36. It has shown that the receptor internalization is often associated with ligand-receptor interaction (30). To understand how hemin modulates the surface CD36 levels, we need to examine the binding environment that facilitates the hemin interaction. To investigate the favourable microenvironment for hemin binding, we have analysed the hemin binding pocket from various hemin/heme co-crystallized proteins and extracted the recurring amino acids and their interaction patterns. The residues aspartic acid (D), phenylalanine (F), methionine (M), arginine (R), serine (S), valine (V) and glutamine (Q) were found to be repeating consistently in seven complexes out of hemin-protein complexes analysed (Table 2). A distance map was generated using hemin and protein atoms, and the distance between atoms was color-coded with the VIBGYOR (violet for the lowest distance to the red for highest distance) (Figure 4). The high resolution atomic-level examination of hemin and its surrounding residues in biophore has revealed that the arginine, aspartic acid and glutamine residue atoms were in close distance with the hemin carboxylate group and provides hydrophilic environment which facilitates the hemin binding (Figure 4).

**Figure 4:**
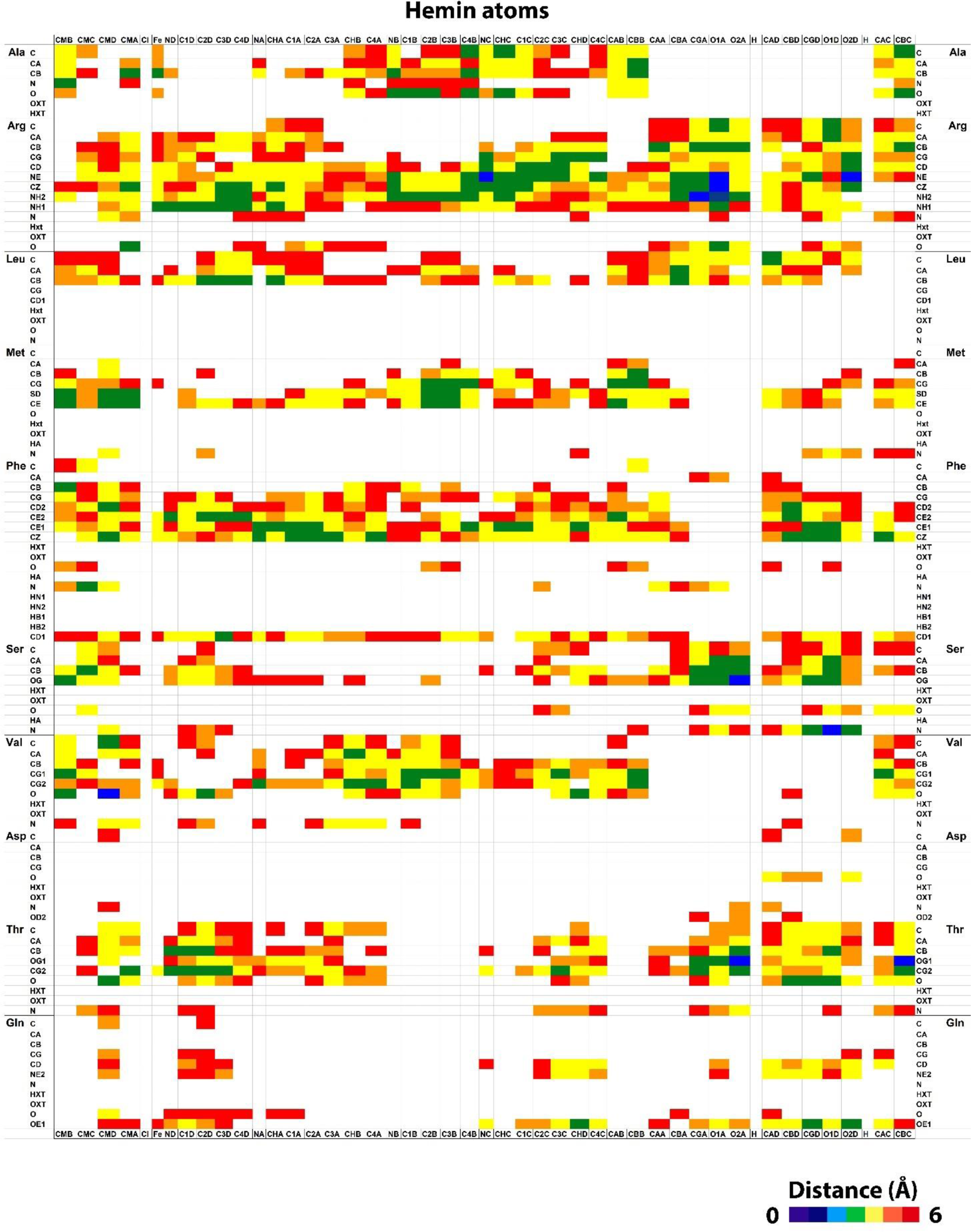
Distance matrix representing hemin biophore. The hemin bound protein complexes were identified and extracted the hemin binding region to ascertain biophoric features. The distance between the hemin atoms to amino acid atoms were analyzed and generated a 105×46 matrix and colour coded with VIBGYOR (violet for lowest distance to red for highest). The matrix represents distance data from a total of eight hemin bound complexes. The carboxylate group of the hemin and the polar side chains of amino acids were found to be enriched in heat map.

**Table 2.** Recurring amino acids in hemin biophore.

| Amino Acid /PDB ID | A | C | D | E | G | H | I | K | L | M | N | P | Q | R | S | T | V | W | Y |
| --- | --- | --- | --- | --- | --- | --- | --- | --- | --- | --- | --- | --- | --- | --- | --- | --- | --- | --- | --- |
| 1O9X | 1 | 0 | 1 | 0 | 1 | 1 | 1 | 1 | 1 | 1 | 0 | 1 | 0 | 1 | 1 | 0 | 1 | 0 | 1 |
| 2BLI | 1 | 0 | 1 | 0 | 0 | 1 | 1 | 1 | 1 | 0 | 0 | 1 | 0 | 1 | 1 | 1 | 1 | 1 | 1 |
| 2QSP | 1 | 0 | 1 | 0 | 0 | 1 | 0 | 1 | 1 | 1 | 1 | 0 | 0 | 0 | 1 | 1 | 1 | 0 | 1 |
| 2R7A | 1 | 0 | 1 | 1 | 1 | 0 | 1 | 0 | 1 | 1 | 1 | 0 | 1 | 1 | 1 | 1 | 1 | 1 | 1 |
| 2ZDO | 1 | 0 | 0 | 1 | 1 | 1 | 1 | 1 | 1 | 1 | 1 | 0 | 0 | 1 | 0 | 1 | 1 | 1 | 0 |
| 3NU1 | 1 | 0 | 1 | 0 | 1 | 1 | 0 | 0 | 1 | 1 | 1 | 0 | 0 | 1 | 1 | 1 | 0 | 0 | 1 |
| 4MF9 | 0 | 0 | 1 | 0 | 1 | 1 | 1 | 1 | 1 | 1 | 0 | 1 | 1 | 1 | 1 | 1 | 1 | 0 | 0 |
| 4MYP | 1 | 0 | 1 | 0 | 0 | 0 | 0 | 1 | 0 | 1 | 1 | 1 | 1 | 1 | 1 | 1 | 1 | 0 | 1 |
|  | 7 | 0 | 7 | 2 | 5 | 6 | 5 | 6 | 7 | 7 | 5 | 4 | 3 | 7 | 7 | 7 | 7 | 3 | 6 |

The relative contribution of each residue in the hemin binding pocket was analysed using webserver ABS-Scan (http://proline.biochem.iisc.ernet.in/abscan/), and their relative binding energies were estimated (Figure 5A). The molecular docking of hemin within CD36 has revealed that the residues R292, Q382 interacting with hemin through hydrogen bonding and D372 residue through salt bridges with the hemin carboxylate group (Figure 5B). The mutation of R292, Q382 to alanine found to be destabilizing the CD36-hemin complex, whereas the mutation in D372 to alanine was found to be increasing the affinity of CD36 towards hemin. Further, the stability of the wild type CD36 or different mutants (R292A, D372A, and Q382A) complexed with hemin, was studied by molecular dynamics simulation using RMSD, radius of gyration and RMSF as evaluating criteria. The average RMSD value of CD36 (wild-type), R292A, D372A and Q382A complexed with hemin were found to be 0.58, 0.82, 1.13 and 0.44 nm respectively. Notably, the D372A mutant showed higher deviation compared to the R292A and Q382A (Figure S4A). The aspartic acid residues contributes to the ionic character of proteins and replacement of it with non-polar alanine residue may imparts hydrophobicity hence higher fluctuation in RMSD (32). The other explanation could be the mutants were constructed on the wild type structure and it may introduced structural perturbations which is reflected by the increased in RMSD (33). The RMSD value of wild type and mutants throughout the simulation suggests the complexes were converged which is an indicator of stability, although the minor perturbations were observed (Figure S4A). The radius of gyration can be used to investigate the compactness of protein-ligand complexes. The average radius of gyration for CD36 (wildtype) was found to be 2.46 nm whereas for R292A, D372A and Q382A it was 2.53, 2.58 and 2.48 nm respectively (Figure S4B). There is no major change in radius of gyration of wild type and mutant complexes indicate the compactness of protein-ligand complexes. Further, the fluctuation of hemin during the simulation was assessed using root mean square fluctuation (RMSF). As it is evident from the (Figure S4C) the fluctuation is minimal among the wild-type and mutants. The overall molecular dynamics studies indicates the stability of CD36 and mutants complexed with hemin.

**Figure 5:**
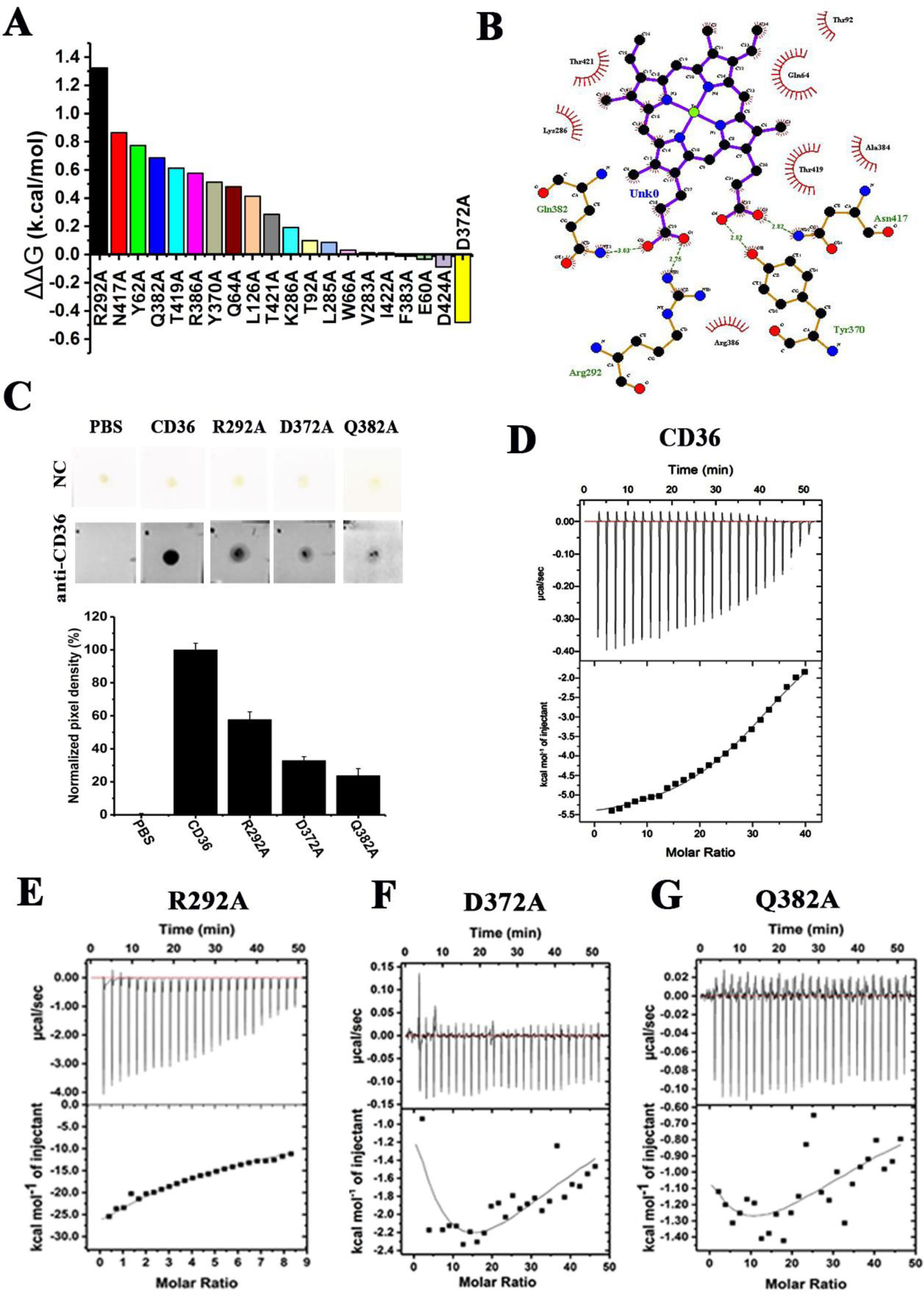
The residues R292, D372 and Q382 are crucial for the active engagement of hemin with CD36. **(A)** The relative contribution of biophoric residues to CD36-hemin affinity. The residues in hemin binding domain on CD36 were analyzed for their contribution towards CD36-hemin interaction using alanine scanning mutagenesis (webserver ABS-Scan). The complex was uploaded into server for predictions. The results were plotted as difference in binding energy of wildtype and mutant. The residues R292, N417, Y62, Q382 were showing significant change in the binding energy indicates they could influence the CD36-hemin interaction. **(B)** The interaction analysis of CD36-hemin complex using ligplot+ software. The residues R292, Q382, Y370 and N417 can be found in interaction through hydrogen bonding (green dotted lines). The other residues K286, T421, T92, Q64, T419 and A384 were found to be associated with hemin through hydrophobic interaction. **(C)** Dot blot assay of CD36 or mutants (R292A, D372A and Q382A) with hemin. After blocking, the hemin spotted individual nitrocellulose membrane strips were incubated with CD36 or mutants for 2 h, washed and probed with anti-CD36 antibodies. The developed blot (anti-CD36 antibody) represents the strong affinity for CD36 and the relative signal intensity quantified is showed in bar graph. A severe reduction in intensity for mutants indicates their prominent role in affinity mediation. **(D)** ITC thermogram of CD36 against hemin. The affinity of hemin towards CD36 was studied using ITC as described in materials and methods. The saturation of the binding indicates all the binding sites were occupied by the hemin. The Kd value for CD36 calculated after fitting the data in binding site model was found to be 1.26±0.24 μM. The mutants R292A **(E)**, D372A **(F)** and Q382A **(G)** were titrated with hemin in identical condition as that of CD36. The Kd values for mutants were calculated and found to be 115±2.14 μM (R292A), 295±5.26 μM (D372A) and 420.16±22.5 (Q382A). The severe reduction in affinity values demonstrate the role of the residues in hemin binding to CD36.

Molecular modelling, molecular dynamics and in-silico mutation studies indicate a suitable hemin biophore and perfect microenvironment to allow the binding of hemin into CD36 ectodomain. To verify these findings, purified CD36ecto wild type and different mutants (R292A, D372A and Q382A) were used in dot blot assay to test their binding towards hemin blotted on the nitrocellulose membrane. The dot blot analysis indicate strong reactivity of CD36ecto wild type with an appearance of intense spot whereas ∼48% signal reduction with R292A and almost 70% reduction in signal for D372A or Q382A (Figure 5C). The affinity of the hemin to the CD36ecto wild type or mutants was investigated using Isothermal calorimetry (ITC). The ITC thermogram of hCD36ecto against hemin indicates the strong binding between the components. The high negative ΔH value suggests the two molecules interacting through hydrogen bonding and van der Waals interactions (Figure 5D). The DP (differential power) change/injection suggests the hemin occupied the binding sites completely to saturate DP values (Figure 5D, upper portion of thermogram). The K_D_ value calculated by fitting the raw data into the two-site binding model and found to be 1.26±0.24 μM and the stoichiometry of hCD36ecto to hemin was found to be 1:2. Similarly, under the identical experimental conditions, the CD36ecto mutants (R292A, D372A and Q382A) were titrated with hemin and the ITC thermogram was recorded. The ITC titration curve for R292A indicated reduction in enthalpy values and increase in entropy values which highlight weak interaction between hemin with CD36ecto R292A. It is probably be mediated by weak non-covalent hydrogen bonding (Figure 5E). The KD value calculated from data fitting was found to be 115±2.14 μM. It is 100 folds lower compared to the wild type CD36ecto. The R292A thermogram clearly indicates that the hemin is unable to fit within binding pocket and as a result it is exhibiting reduced affinity towards hemin (Figure 5E). The thermodynamic parameters of R292A is purely due to the disruption of hemin favourable biophore microenvironment. The ITC thermogram of other two mutants (D372A or Q382A) revealed that the mutation has drastically reduced the affinity of hemin towards CD36 ectodomain (Figure 5F and Figure 5G). The Kd value calculated for D372A or Q382A mutants were found to be 295 ± 5.26 μM and 420.16 ± 22.5 respectively. The significantly higher Kd values for D372A and Q382A suggests the crucial role of these residues in hemin binding pocket. To rule out the observed effects are not due to the structural changes or aggregation of mutant proteins, the circular dichroism spectrum and size exclusion chromatography was performed on purified mutant proteins. The size exclusion elution profile and CD spectrum analysis indicates there was no change in oligomer status and secondary structural element in mutant protein (Figure S5A and S5B). The cumulative results represented in Figure 5, suggest that the residues R292, D372 and Q382 are crucial for hemin binding and there are sufficient evidences to claim that hemin act as a ligand for CD36.

### CD36-hemin interaction on cell surface is crucial for immune-dysfunction in macrophages

The scavenger receptor CD36 acts as a receptor for various ligands from diversified sources at cellular level (34). It has been shown that CD36 also acts as a co-receptor for TLRs which are also involved in innate-immune response (35). As professional macrophages J774A.1 harbours functional CD36 and TLRs on their cell surface to contribute into hemin mediated innate immune response. To understand the role of CD36 in hemin mediated immune response, we have screened several cell line with a requisite phenotype of TLR^-/-^ and CD36^-/-^ (Figure S6A, S6B and S6C). The flow cytometry based screening suggested that the MG63 cells have no or very low levels of CD36 level and TLR expression on their cell surface (36, 37). The cells transfected with wild type CD36 showed enhanced migration towards the hemin placed in lower chamber whereas very few number of cells were migrated in absence of hemin (Figure 6A, wild type panel). The R292A transfected MG63 cells showed migration in presence of hemin but in case of D372A or Q382A transfected cells, the migration is very less (Figure 6A). The chemotaxis index for CD36 transfected cells in presence of hemin was found to be 31 folds compared to the non-specific migration (chemophoresis) in absence of hemin (Figure 6B). Disruption of hemin biophore in CD36 through mutation in ectodomain abolishes the chemotaxsis towards hemin but didn’t affect the non-specific migration of cells (Figure 6B). The MG63 transfected with empty vector (mock transfected) didn’t allow chemotaxis towards hemin further strengthen the observation that functionally active CD36 on the cell surface is crucial for chemotactic migration of cells towards hemin. These experimentation results provide direct evidences that hemin is being recognized by CD36 presented on cell surface with high affinity and specificity. Now we explored whether the CD36-hemin interaction is responsible for immunological responses in macrophages under malaria like environment. The MG63 cells with wild type CD36 is exhibiting enhanced phagocytosis in response to hemin (25 µM) treatment (Figure 6C). In comparison to wild type, phagocytosis index of the MG63 transfected with mutant R292A, D372A or Q382A were not showing enhanced phagocytosis with hemin at 25µM (Figure 6C). The phagocytic pattern of mutants R292A, D372A and Q382A clearly highlights role of CD36-hemin interaction in dysregulation of phagocytic activity of macrophages (Figure S7A). Similar to phagocytosis, MG63 transfected with wild type CD36 was exhibiting reduction in bactericidal activity in presence of hemin (0, 25 or 200µM) whereas MG63 cells transfected with different mutants (R292A, D372A and Q382A) were showing no significant effect of hemin exposure. (Figure 6D and Figure S7B). Presence of functional CD36 on cell surface to interact with hemin is crucial for observed functional defects in macrophages.

**Figure 6:**
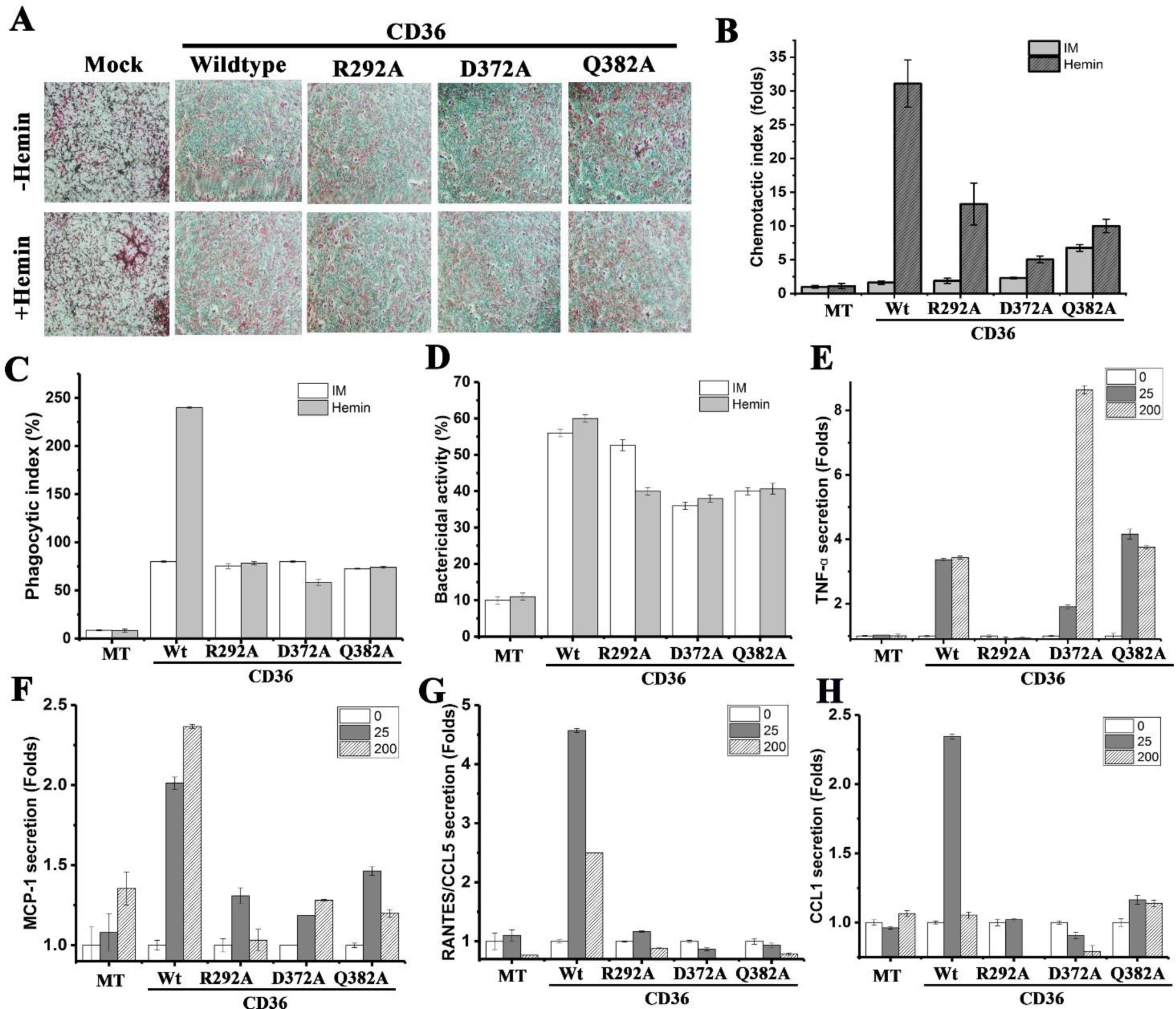
The CD36 is the key receptor for the hemin at cellular level and responsible for the immune-dysfunction. **(A)** Hemin interacts with CD36 at cellular levels. The MG63 cells transfected with either CD36 or mutants (R292A, D372A and Q382A) and carried out migration assay in presence or absence of hemin. The number of cells migrated in CD36 transfected MG63 in presence of hemin is significant and indicates the hemin interacts with the cell membrane bound scavenger receptor CD36. (n=3). **(B)** The images captured after cell migration in CD36 or mutants transfected cells in presence or absence of hemin. The upper panel shows absence of hemin and the lower panel for cells migrated in presence of hemin. There is no clear distinction was observed in mutants or mock transfected cells either in presence or absence of hemin. **(C)** Phagocytic activity of CD36 or mutants transfected MG63 cells. The cell after transfection was either treated with hemin or remain untreated and performed the phagocytosis assay as described in materials and methods section. The results clearly indicates at 25 μM hemin the phagocytic index increased to 30% compared to untreated and CD36 transfected cells. There is no significant change was observed in mutants transfected **25** μM hemin treated cells. **(D)** Bactericidal activity of CD36 or mutant transfected MG63 cells. There is no considerable changes were observed in bactericidal activity either hemin treated or untreated transfected cells. The MG63 cells transfected with CD36 or mutants and treated with hemin (25 or 200 μM). The cytokines TNF-α **(E)**, RANTES/CCL5 **(F)**, MCP-1 **(G)**, and CCL-1 **(H)** were quantified and expressed in folds change. All four cytokines were upregulated to several folds in 25 μM hemin treatment whereas two cytokines RANTES and CCL1 were reduced in 200 μM hemin treated condition.

### Scavenger receptor CD36 is crucial for hemin mediated inflammatory response

It has been reported that the scavenger receptor CD36 interaction with its ligands (endogenous or exogenous) primes the cytokine signalling and could be responsible for host pathology (38). The cytokine array revealed that several pro-inflammatory cytokines up-regulating upon hemin stimulus [Figure 3A, B]. We have further explored the CD36-hemin interaction as an axis to regulate the cytokine secretion. MG63 cells transfected with wild type CD36 or different mutants (R292A, D372A and Q382A) were either remain untreated or treated with hemin (25 or 200µM) for 1hr and secreted cytokines were measured as described in material and methods. The pro-inflammatory cytokines TNF-α (39), MCP-1 (40), RANTES (41), and CCL1 (42) were directly linked to the pathology associated with severe malaria. The cytokine TNF-α produced by the macrophages during various infectious or non-infectious diseases, and responsible for regulating the pathophysiology of the host (43). The TNF-α levels in MG63 cells transfected with wild type CD36 was found to be 4.5 folds higher with hemin treatment compared to untreated cells (Figure 6E). In mock or R292A transfected MG63 cells, TNF-α levels were reduced significantly and it is in agreement with our chemotaxis results. The MG63 cells transfected with D372A showing significantly high level of TNF-α (8.7 folds) at 200 µM hemin compared to MG63 cells transfected with wild type CD36 (Figure 6E). Contrary to other mutants, Q382A is giving very high TNF-α with an identical pattern to the wild type CD36 (Figure 6E). The monocyte chemo-attractant protein 1 (MCP1) plays crucial role in migration of monocytes/macrophages (44). The wild type CD36 transfected (untreated) cells showed basal levels of MCP-1 but secretion went up 2 folds after treatment with hemin (Figure 6F). In R292A mutant transfected cells, MCP-1 secretion was much reduced and secretion pattern is different from wild type CD36 transfected cells. It is exhibiting complete suppression of secretion at high hemin (200µM) treatment. D372A transfected MG63 cells were exhibiting similar secretion pattern as observed for wild type CD36 transfected cells. MCP-1 secretion from Q382A transfected MG63 cells was much low but it follows similar secretion pattern as observed for wild type CD36 transfected cells (Figure 6F). The cytokines RANTES and CCL1 were found to play crucial role in cerebral malaria pathology (45). The RANTES levels in CD36 transfected MG63 cells were found to be increased to 4.6 folds at low hemin (25µM) but it was reduced significantly at 200 µM hemin (Figure 6G). MG63 cells transfected with mutants (R292A, D372A and Q382A) was exhibiting basal level of cytokine secretion and cells were not showing any up-regulation in cytokine secretion in response to hemin stimulation (Figure 6G). The CCL1 levels in wild type CD36 transfected MG63 cells were found to be upregulated two folds at low hemin (25µM) but it was reduced back to basal level at 200 µM hemin (Figure 6H). MG63 cells transfected with mutants (R292A, D372A and Q382A) was exhibiting basal level of cytokine secretion and cells were not showing any up-regulation in cytokine secretion in response to hemin stimulation (Figure 6H). The data presented in figure 6 further confirm the role of CD36-hemin interaction and its crucial role in functional defects and immune responses in macrophages. More-importantly, these effects are independent to TLR and purely mediated by CD36 and its ability to bind hemin in macrophages.

### Hemin induces the phosphorylation of membrane bound CD36

The phosphorylation of proteins during intracellular signalling is crucial as it controls various cellular events (46). Since the hemin modulating the immune-response through scavenger receptor CD36 we are interested to know the signal transduction process associated with CD36-hemin interaction. To determine whether hemin binding CD36 is causing phosphorylation of membrane bound receptor. We have treated the cells with different concentration of hemin (0, 25µM, 200µM) for 1 hr and immune-precipitated the CD36 from untreated or hemin treated lysates. The IP elute was resolved on SDS-PAGE and probed with anti-phosphotyrosine antibody. Hemin treatment leads to the phosphorylation of membrane bound as well as soluble CD36 present in cytosol (Figure 7A). The CD36 phosphorylation follows biphasic pattern with different concentration of hemin (low/high). The cells treated with low concentration of hemin (25µM) is exhibiting 40% more phosphorylation of CD36 present on plasma membrane whereas 10% soluble CD36 was phosphorylated in cytosol (Figure 7B). Cells treated with high hemin (200µM) is showing inhibition of CD36 phosphorylation with appearance of very low level of phosphorylated CD36 on membrane and soluble form in cytosol (Figure 7B). CD36 phosphorylation in macrophages in response to different concentration of hemin (low/high) might explain the immunological responses from macrophages.

**Figure 7:**
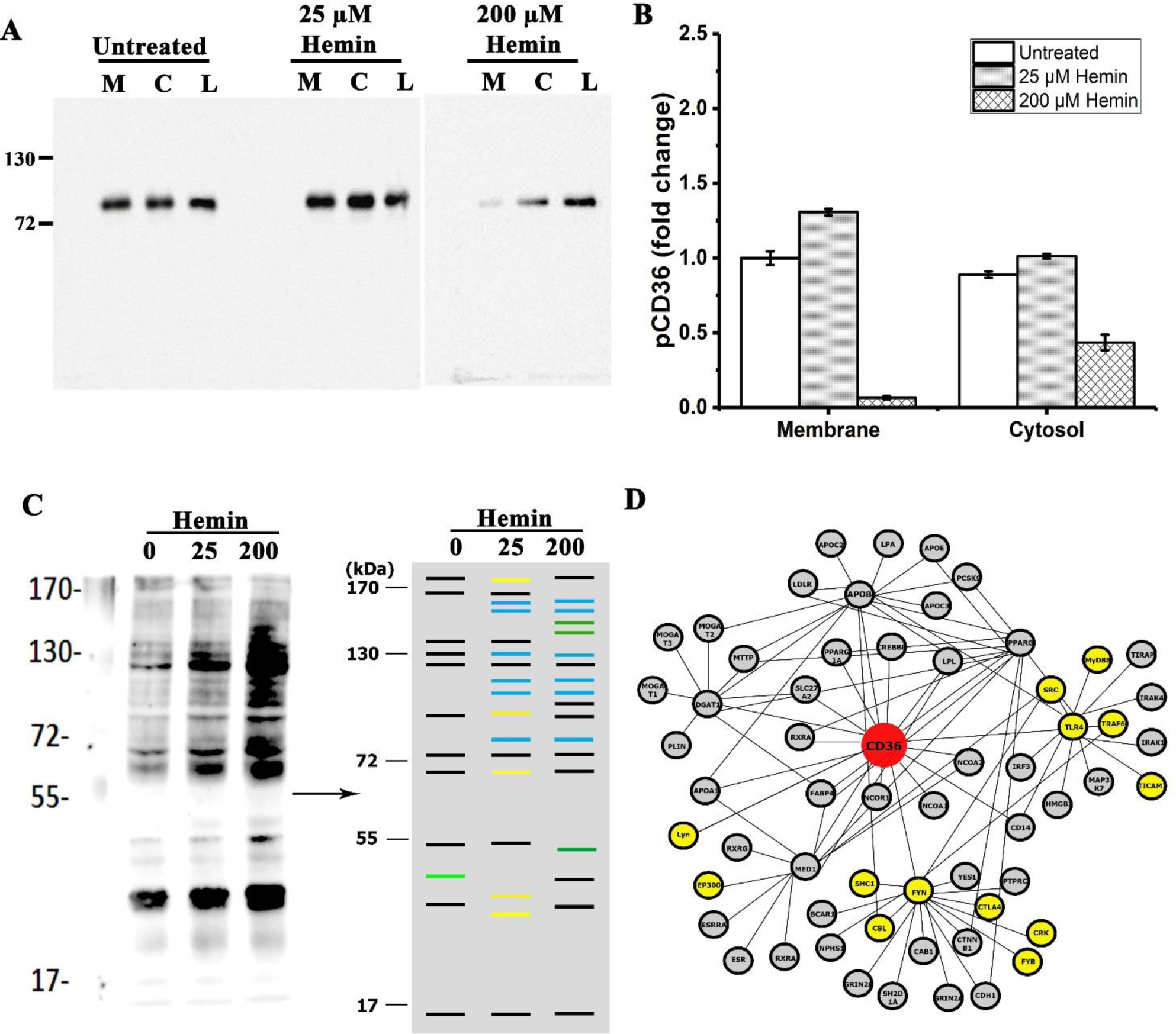
Hemin treatment induces the phosphorylation of CD36. **(A)** The cells either hemin (25 or 200 μM) treated for 1 h or remain untreated were lysed and fractionated into membrane and cytosol. The lysated immunoprecipitated with anti-CD36 antibody as described in materials and methods section. The resolved proteins transferred onto membrane and probed with anti-phosphotyrosine antibodies. The enhanced phosphorylation of CD36 (cytsolic or membrane fractions) in 25 μM hemin treated cell lysates can be observed whereas in 200 μM hemin terated cells the reduction of phosphorylation was observed. The blots were analyzed for signal intensity and the fold change in the CD36 phosphorylation was plotted in **(B). (C)** The cell lysates prepared from hemin treated or untreated macrophages, resolved on SDS-PAGE and transferred onto membrane. The membrane was probed with anti-phosphotyrosine antibody indicates the activation of several proteins in hemin treated condition. A mock blot was developed to clearly represent the differential phosphrylation of proteins from untreated to treated condition. The proteins exclusively phosphorylating in 25 μM hemin treated lysates coloured in Yellow and the phosphoproteins exclusive to 200 μM color coded in dark green. The phosphoreactive bands common in 25 and 200 μM lanes color coded in Blue. **(D)** Webserver Cluspro predicted adaptor proteins downstream to CD36. To predict possible adaptor protein downstream to CD36 a protein-protein docking was carried out using the C-terminal domain of CD36 as a bait. The interactome indicates the association of Src family kinases to CD36.

### CD36-Hemin interaction is relaying down-stream signalling

The enhanced phosphorylation of CD36 in hemin treated macrophages indicates the possible signalling event associated with CD36-hemin interaction. We have treated the cells with different concentration of hemin (0, 25µM, 200µM) for 1hr and cell lysate was probed with anti-phospho tyrosine. Further, the analysis of molecular weight of phosphoreactive bands suggested that various proteins with molecular weight between 50-100 kDa were found to be phosphorylating (Table S2). The prediction of different phosphor-protein using phosphoprotein database (https://www.phosphosite.org/homeAction.action) gives list of potential candidate present in different treatment groups. Macrophage treated with hemin (low/high) is causing appearance of Nitric oxide synthase (Nos1), Receptor-type tyrosine kinase FLT 3 (FLT3), E3 Ubiquitin ligase (CBL), serine threonine protein kinase D1 (PRKD1) compared to the untreated cells. Whereas, macrophage treated with low hemin (25µM) is causing appearance of RB1-inducible coiled-coil protein 1 (RB1cc1), STAT1, STAT3, proto-oncogene Src, Yes, Fyn and Heat shock protein β-1 (HSP27). It is also showing phosphorylation of CD36 as well (Figure 7C). Interestingly, few exclusive phosphorylated protein bands appeared in macrophage treated with high hemin (200µM). Ribosome binding protein 1 (RBP1), Mitogen-activated protein kinase kinase kinase kinase 4 (MAP4K4), Adenylate cyclase type 8 (Adcy8) and protein C-ets-2 (ETS-2) was phosphoprotein appeared in macrophage treated high hemin (200µM) compared to untreated cells (Figure 7C). CD36 phosphorylation recruit several adaptor proteins to relay intracellular signalling for different immune responses (47). To explore these adaptor proteins down-stream to CD36-hemin signalling, we have used C-terminal end of CD36 as a bait protein to identify the interacting partner using Cluspro web server (https://cluspro.bu.edu/). The server is predicting the binding efficiency of different adaptors with the bait protein and provide binding energy for the resulting complexes (supplementary Table S3). The results suggest the top hits could be from NF-κB/STAT signalling, Ras kinase signalling, AKT1, MAPK1/ERK2, STAT signalling pathway (Table S2). The predicted adaptor protein interactome of CD36 is presented in (Figure 7D).

### Tyrosine kinase Lyn is down-stream to the CD36-hemin signalling axis

Src family tyrosine kinases were reported to down-stream to the receptor mediated down-stream signalling in response to extracellular ligands (48). Several potential adaptor proteins from src family were predicted by Cluspro web server (https://cluspro.bu.edu/). We have treated J774A.1 with either remain untreated or treated with hemin (25µM), CD36 was immunoprecipitated and re-probed with respective src family kinases antibodies (Lyn, Fyn, Src, Lck, Csk) as described in material and methods. The immunoblot analysis with anti-Yes indicate that the Yes is associated with CD36 and hemin treatment is not changing the level of protein associated with CD36 (Figure 8A). It indicates that the Yes has strong affinity to the membrane bound CD36 and confirm our earlier prediction of the Cluspro web server. C-terminal src kinase and src kinase have very low affinity to CD36 and the protein levels remained unchanged (Figure 8A). Probing the blot with anti-Fyn is not giving any band with untreated but give cryptic band in hemin treated cells but there is no change in protein level associated with CD36 (Figure 8A). Testing the presence of Lyn in the IP eluent, we have found change in the level of protein associated with CD36. A very little protein was associated with CD36 in untreated cells compared to high amount in hemin treated J774A.1 cells (Figure 8A). To confirm the interaction of Lyn with CD36, we have immune-precipitated Lyn from untreated or hemin treated cells and probed the IP eluent with anti-CD36 antibodies. As expected, there was a change of level of CD36 between untreated and hemin treated cells. A very little CD36 was present in the untreated cells compared to high amount in hemin treated J774A.1 cells (Figure 8B). It has been shown by various research groups that CD36 downstream association is often found with src family adaptor proteins (49). The recruitment and activation of Lyn to the docking site leads to the activation of Lyn and recruitment of Vav and FAK proteins which initiates the macrophage foam cell formation (50).

**Figure 8:**
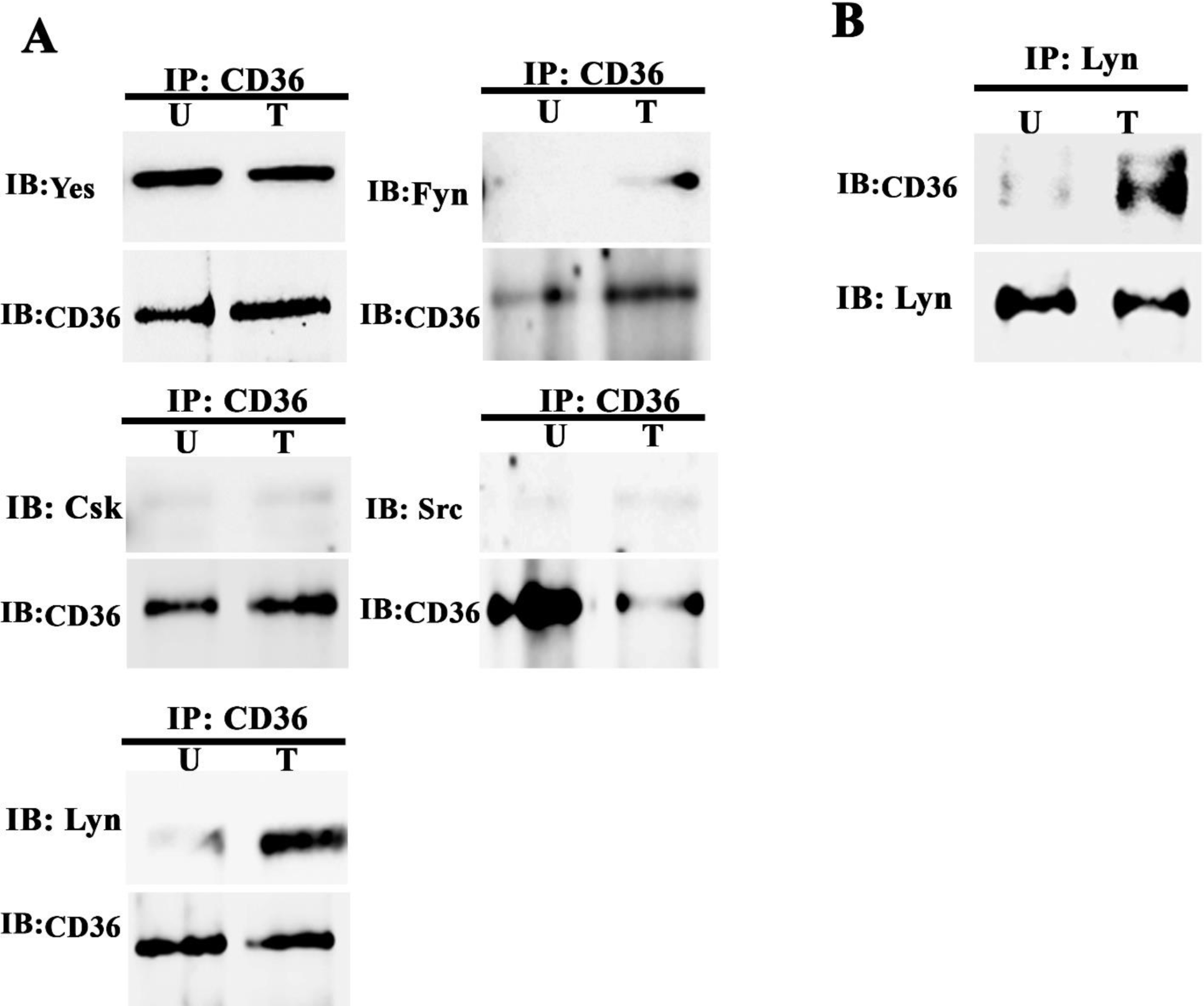
CD36 associates with Src family kinases in hemin treated macrophages. **(A)** The macrophages either treated with hemin for 1 h or remain untreated were lysed in IP lysis buffer supplemented with protease inhibitors. The cell lysates immuno-precipitated with anti-CD36 antibody as described in methodology and probed with anti-Yes, anti-Fyn, anti-Csk, anti-Src and anti-Lyn antibodies or anti-CD36 antibody. A significant co-precipitation of Lyn with CD36 was observed in hemin treated macrophage cell lysates. Besides a minor levels of Fyn kinase found to be associated with CD36 in hemin treated cells. More-over, the Yes, Csk and Src kinases were found to be constitutively associated with CD36. **(B)** The macrophages cell lysates from treated with hemin or left untreated condition, immuno-precipitated with anti-Lyn antibody and probed with anti-CD36 or anti-Lyn. The (A) and (B) confirms the recruitment of Src family kinase Lyn to downstream to CD36 in hemin treated condition.

### Tyrosine kinase Lyn is controlling immune responses down-stream to the CD36-hemin signalling

The main complication with hemin treated macrophages is the secretion of pro-inflammatory cytokine especially TNF-α. Lyn is present as an adaptor protein down-stream to the CD36-hemin signalling. Now, we have explored whether Lyn has a role in cytokine secretion and immune responses from macrophages treated with hemin. Lyn was knockdown in J774A.1 using siRNA and level of lyn was confirmed by western blotting using anti-Lyn antibodies (Figure 9A). Lyn knockdown in J774A.1 is reducing the secretion of TNF-α compared to the scrambled siRNA treated control cells. There is almost more than 50% reduction in the level of cytokine in Lyn knockdown J774A.1 cells (Figure 9B). J774A.1 contains toll-like receptors and other hemin responsive receptors, there is a possibility that remaining cytokine secretion could be due to the hemin interaction with these receptors (51).

**Figure 9:**
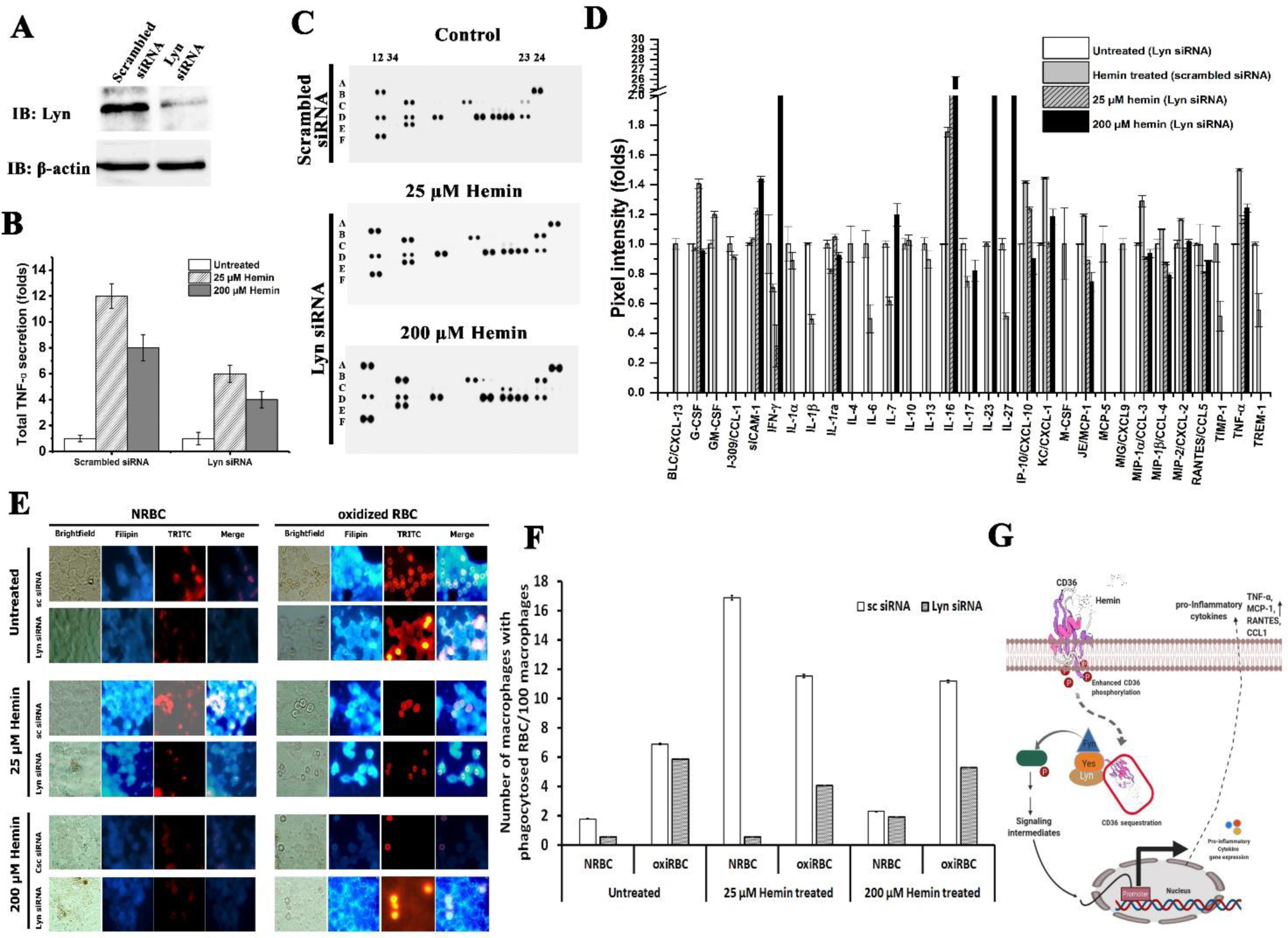
Recruitment and docking of Src family kinase Lyn to downstream of CD36 is crucial for hemin mediated immune-dysfunction. **(A)** The validation of Lyn knockdown in macrophages. The macrophages either transfected with scrambled siRNA (sc siRNA) or Lyn siRNA and 48 h post transfection, lysates were probed with anti-Lyn antibody or anti-β actin antibody. The blot shows the 70-80% knockdown of Lyn protein. **(B)** Estimation of TNF-α levels from Lyn knockdown macrophage cell culture supernatants. The lyn knockdown macrophages treated with hemin showed reduced levels of TNF-α comparing to scrambled siRNA transfected macrophages indicates the hemin activates the cytokine signalling through CD36 and recruitment of lyn to the cytosolic domain of CD36. **(C)** Global cytokine profiling of Lyn knockdown macrophages. The macrophages were transfected with scrambled siRNA or Lyn target siRNA and either untreated or treated with hemin. The cytokine array was carried out on cell culture supernatants using mouse cytokine profiler array kit panel A. **(D)** The cytokines were normalized and plotted as fold change. Several cytokines such as CXCL-10, CXCL-1, MCP-1, CCL-3, CCL-4, CCL-2, and TNF-α levels were reduced (expressed in fold change) upon lyn knockdown. **(E)** The phagocytic activity of macrophages (with lyn knockdown) towards RBC. The macrophages either transfected with scrambled siRNA (sc siRNA) or or Lyn siRNA were either treated with hemin or remain untreated were incubated with normal RBC or oxidized RBC. Macrophages were stained with filipin to identify phagosomes containing engulfed material and observed under Nikon 80i fluorescence microscope in DAPI and TRITC filters to acquire filipin (Blue) and RBC (Red) fluorescence respectively. **(F)** The number of phagocytic macrophages were counted and plotted as number of phagocytic macrophages per 100 counted macrophages. The lyn knockdown in macrophages significantly reduced the non-opsonic phagocytosis of NRBC or oxiRBC in hemin treated condition. **(G)** Proposed hypothesis for hemin activated CD36 downstream signalling. The hemin interaction with CD36 induces the CD36 phosphorylation and sequestration of CD36 into intracellular vesicles. Further, the hemin binding to CD36 activates the intracellular signaling by recruitng the adaptor proteins from Src family to the cytoslic domain of CD36. The CD36-adaptor protein signalling complex intiates the cascade of events that trigger the transcription of pro-inflammatory cytokine genes.

The global cytokine profiling in absence of Lyn in hemin treated macrophages were studied using cytokine array. Gene silencing of Lyn in macrophages was found to down-regulate CXCL-13, GM-CSF, CCL-1, IL-1α, IL-1β, IL-4 and 6, M-CSF, CXCL-9, MCP-5 and TIMP-1 in response to hemin simulation (Figure 9C and 9D). Interestingly, Lyn silencing in J774A.1 is up-regulating IL-16 levels 26 folds in response to hemin (25 or 200µM) stimulation. In addition, cytokine IL-23, IL-27 were found to up-regulated in Lyn knockdown macrophages in response to high hemin (200µM) but absent in low hemin (25µM). The pro-inflammatory cytokine IP-10 directs homing of T-cells and promote inflammation in the cerebral region to contribute into cerebral malaria pathology (52). The elevated levels of IP-10 is associated with cerebral malaria development in mouse models (53). The macrophage stimulated with hemin (25 µM) and transfected with scrambled siRNA is exhibiting 1.41 folds enhancement in IP-10 levels. Whereas Lyn knockdown in macrophages is reducing to normal level in response to hemin stimulation. (Figure 9C and 9D). This might explain the elevated levels of IP-10 due to extracellular hemin correlate with cerebral malaria pathology (54). The dysregulation of TNF-α level is associated with several infectious and non-infectious diseases. The TNF-α levels during cerebral malaria is linked to very high parasite load and anaemia (55). Knock down of Lyn in macrophages was found to down-regulate TNF-α in response to hemin simulation (Figure 9C and 9D). The global cytokine profiling in J774A.1 confirms the hemin mediated inflammatory response requires recruitment of Lyn to CD36 downstream.

Further, we have evaluated the phagocytic activity in Lyn knockdown macrophages using normal and oxidized RBC as phagocytic objects (Figure 9E). The control cells treated with 25 μM hemin showed phagocytosis of normal as well as oxidized RBC. Conversely the higher hemin concentration (200 μM) abolished the erythrophagocytosis. Interestingly, the Lyn knockdown in J774A.1 treated with 25 μM or 200 μM showed phagocytosis of oxidized RBC but not normal RBC (Figure 9F). Moreover, the Lyn knockdown does not affected the hemin mediated phagocytic depression and bactericidal activity towards bacteria (Figure S8A and S8B). The selective phagocytosis of oxiRBC is controlled by CD36 and its downstream adaptor Lyn in hemin treated macrophages. The results strongly suggest that hemin binding to CD36 causes the CD36 phosphorylation at tyrosine residue and recruitment of signalling protein Lyn kinase to the cytosolic domain. Further, the hemin-CD36-Lyn involved in the pro-inflammatory cytokine secretion (Figure 9G).

## Discussion

Macrophages in host body serves as the first line of defence against pathogens or endogenous PAMPS (56). The macrophages treated with PAMPS or endogenous ligands secrete pro-inflammatory or anti-inflammatory cytokines such as TNF-α, IL-1*β,* and Interleukins (57). During malaria, macrophages are activated by sensing of IRBC and the parasite proteins like PfEMP-1 (58). The macrophages phagocytose the IRBCs and induce innate immune response by secreting IL-10, IL-12, IL-18, TNF-α, IFN-γ cytokines and generate oxidative stress (59). The most prominent pro-inflammatory cytokines associated with malaria infection are IFN-γ, TNF-α, MCP-1, IL-6, IL-8, RANTES, IP-10, and IL-18 (60). In placental malaria, the macrophages cause local inflammation and low birth weight by releasing the cytokines after activation signal from sequestered IRBCs from placenta. Macrophage receptors also facilitate the non-opsonic phagocytosis of IRBC. The expression of PS on cell surface is essential for the phagocytosis by macrophages. During vascular injury or oxidative stress the PS exposed on RBC surface (61). The macrophage uses various cell surface receptor to sense the senescence or IRBCs (62). The prominent macrophage receptor that recognizes the IRBC is scavenger receptor CD36. The CD36 not only recognizes PS expressing RBC (14) but also serves as a receptor for parasite PfEMP-1 (63). Several line of evidence highlighted the CD36 mediated non-opsonic phagocytosis is crucial to check the unbalanced pro-inflammatory cytokine secretion.

The hemin (free heme) released during vascular injury, or malaria implicated in pathological outcomes of host (54). The hemin is highly cytotoxic and removed by host machinery through hemopexin and further catabolized by heme oxygenase-1 (64). During hematoma a large amounts of hemin (10mM) released in brain and could lead to stroke and permanent damage to brain (65). The cytotoxicity of heme is attributed to the hydrophobic porphyrin ring which can easily intercalates into plasma membranes and damage the cell integrity and production of free radicals (66). The exposure of RBC to hemin causes PS exposure on RBC surface. The PS expressing RBC removed from circulation by non-opsonic phagocytosis by macrophages and cause anaemia. Intravascular haemolysis during malaria releases large amounts of free heme which can be deposited in liver. The free heme activates NF-κB which induces the VCAM-1, KC and MIP-1 cell adhesion molecules and cytokines expression (67). The heme treatment induced TNF-α secretion by macrophages through TLR-4, CD18 and MyD88 dependent manner (68). The expression of cytokines attract the neutrophils and their attachment to liver tissue. The levels of free heme in liver is correlating with the neutrophil extravasation and liver damage (69).

The scavenger receptor CD36 play crucial role in controlling the pathophysiology of the host during infectious and non-infectious diseases (70). The macrophages expressing CD36 recognizes various ligands from pathogens or host cells, get activated and secret pro-inflammatory cytokines (71). The binding of oxiLDL with CD36 triggers the association of CD36 with Toll like receptors (TLR-4 and 6) and responsible for sterile inflammation in macrophages (72). The macrophage CD36 mediated oxiLDL internalization activates pro-inflammatory cytokines, generation of ROS which is involved in macrophage foam cell formation in atherosclerosis (73). The binding of TSP-1 to endothelial CD36 leads to apoptosis of endothelial cells (74). The scavenger receptor CD36 mediated pro-inflammatory cytokine secretion is also attributed to ischemic brain injury. Further the CD36 inhibition in mouse ovarian cancer xenografts, reduced the tumour burden, and intracellular oxidative stress (75). The CD36 knockdown reduced the pro-inflammatory cytokine secretion, macrophage migration, ROS generation and plaque formation (76). The macrophages with CD36 knockdown are not efficient to perform non-opsonic phagocytosis and fatty acid uptake which might affect the functioning of the organs (77, 78). The reduction in CD36 levels during tuberculosis provided protection against tuberculosis (79). The macrophages exposed to hemin showed phagocytic depression, reduced bactericidal activity, and marked increase in levels of pro-inflammatory cytokines especially TNF-α. Hemin has been reported to induce pro-inflammatory cytokine TNF-α in macrophages (80). Moreover the CD36 is translocated to cytosol in hemin treated macrophages. The ligand binding to CD36 often involved internalization of ligand-CD36 complex (30). Earlier it was reported the heme interact with pattern recognition receptor TLR-4 but the mechanistic details elusive (81). Our study findings suggests hemin interact with scavenger receptor CD36 in a well-defined region consisting of R292, Q382 and D372 residues. This explains the observed CD36 translocation into cytosol hemin treated macrophages. The CD36 co-operation with TLRs is crucial for phagocytosis of pathogens and induce pro-inflammatory cytokine secretion (35). Contrary to the previous observation, we have found that CD36 solely interact with hemin as evident from chemotaxis assay in MG63 cell line. Our observation further supported by CD36 cooperation with TLRs only required for enhanced phagocytosis of IRBC (35). Further, the evaluation of phagocytosis, bactericidal activity and cytokine secretion of MG63 cells transfected with wild-type (CD36) or mutants (R292A, D372A and Q382A) further confirmed that the hemin interactions with CD36 is highly specific and need intact biophore. The study also establishes CD36 as an independent receptor for hemin. However, the TLRs shown to be a primary receptor for hemin, it is worth investigating whether in presence of TLRs, still CD36 accept hemin as a ligand. During ischemic injury heme activates CD36-TLR1/2 signalling and association of MyD88 downstream to TLR1/2 (82). The phospho CD36 levels were enhanced in membrane and cytosol fractions of hemin treated macrophages. It could be possible that the phosphorylated CD36 gets internalized into intracellular vesicles and there is impairment in cellular machinery to allow recycling of receptor back to cell surface. However, a mechanistic study need to be warranted to explore the CD36 trafficking in cytosol and its inability to recycle back. The engagement of ligand to membrane bound receptors has been shown to activate intracellular signalling and phosphorylation of signalling molecules (83). Moreover, the adaptor protein screening revealed possible adaptor molecules from NF-κB/STAT signalling, Ras kinase signalling, AKT1, MAPK1/ERK2, STAT signalling pathway. Previous studies suggested that immediate downstream signalling molecules down-stream to CD36 are mostly belongs to Src family kinases (84). The pull-down assay using anti-CD36 antibodies suggested the recruitment of Src family kinase proteins Lyn and Fyn to the cytosolic domain of CD36 in hemin treated macrophage cell lysates whereas the Yes, Csk and Src found constitutively associated with CD36. The CD36-dependent signalling cascade requires the recruitment of src kinases Fyn and Lyn, and the MAP kinase kinase and the JNK2 (85). The heme binding activates TLR mediated downstream signalling involving recruitment of TIRAP and MyD88 to the cytosolic side of TLRs. The intermediate signalling molecules TRAF6 and TAK1 activates the NF-κB transcription which induce pro-inflammatory cytokine secretion (86). There is a possibility that heme mediated downstream signalling is receptor specific rather than generalized. Further, the Lyn knockdown macrophages treated with hemin showed reduced pro-inflammatory cytokine secretion. Further, the lyn knockdown macrophages showed improved phagocytosis of oxiRBC but not the bacteria suggests the CD36 is specifically phagocytose oxiRBC through non-opsonic phagocytic mechanism and requires the recruitment of Src family kinase adaptor Lyn to downstream. Although we have predicted adaptor molecules to downstream to CD36-hemin complex, the intermediate adaptor molecules and transcription regulators have not been explored in this study. The identification of complete signalosome may provide more precise information regarding the pro-inflammatory cytokine signalling. Our study provides direct evidence that hemin act as a ligand for CD36 and likely to contribute the pathology of cerebral malaria through CD36.

## Supporting Figures

**Figure S1:**
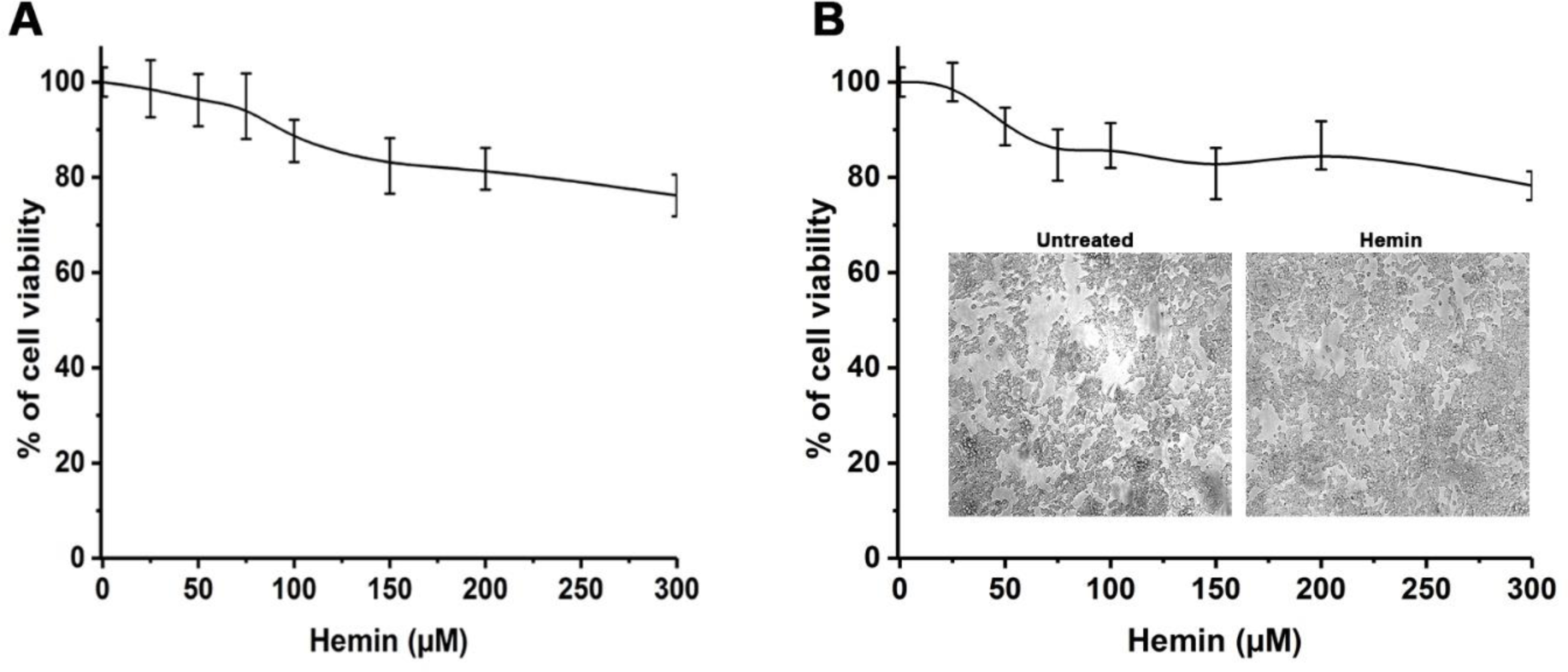
Hemin do not reduce the cell viability of macrophages. **(A)** The macrophages were treated with hemin dose dependently (0 to 300 µM) in presence of serum for 1 h and the cell viability assessed by MTT assay. **(B)** Similarly the cell viability of macrophages assessed in serum free medium. The cell viability in both the conditions suggests the hemin even at higher concentration (200 µM) is not affect the macrophage cell viability.

**Figure S2:**
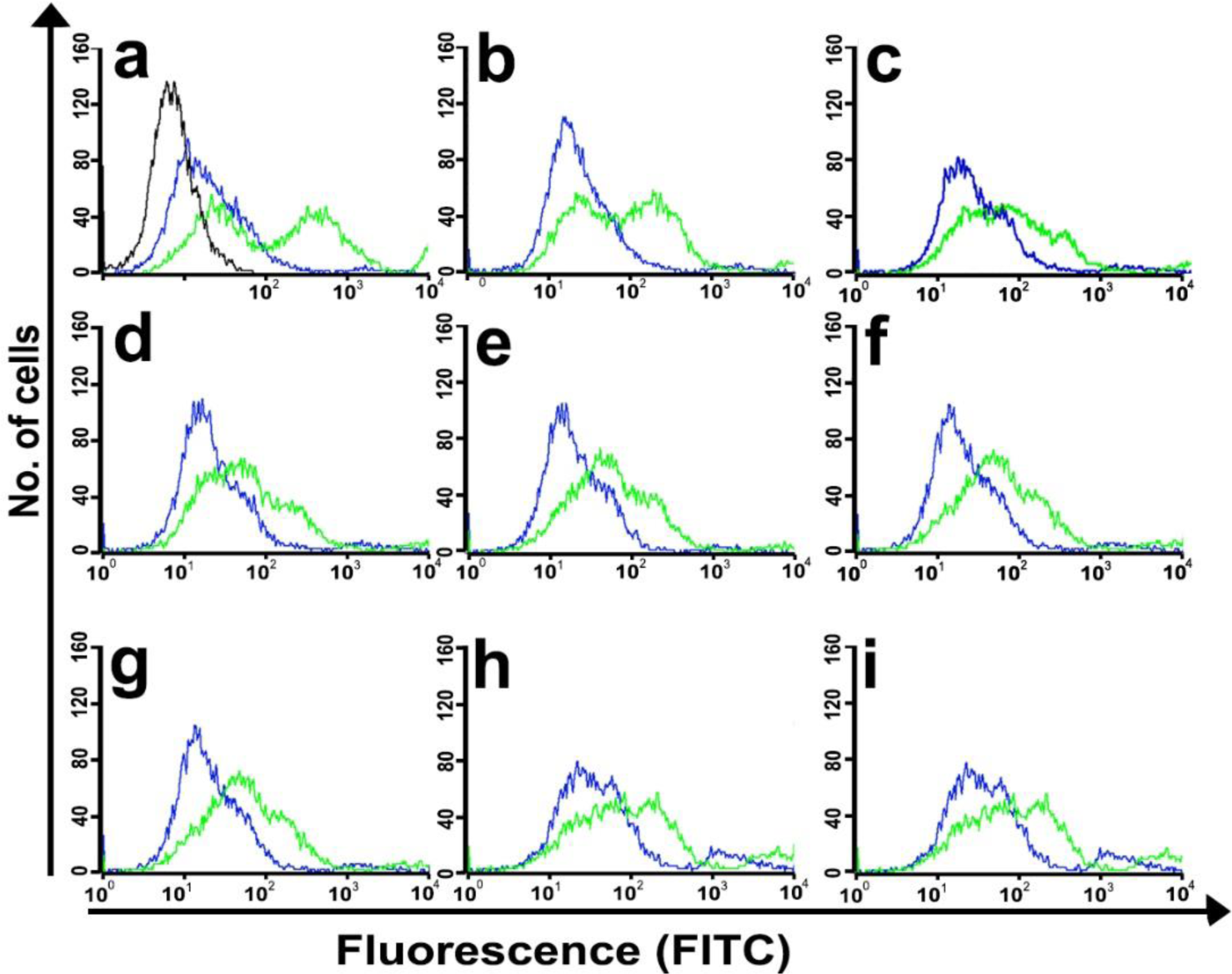
Evaluation of phagocytic activity of macrophages in a flow-based assay. Measurement of Phagocytosis of hemin treated macrophages in flow cytometry-based assay. Macrophages in different Panels; (a) Untreated (b) 25µM (c) 50 µM (d) 75 µM (e) 100 µM (f) 125 µM (g) 150 µM (h) 175 µM (i) 200 µM hemin treated macrophages. Green colour curve is macrophage allowed to phagocytose FITC labelled bacteria whereas blue coloured curve after adding 1% trypan blue. Black coloured curve in panel “a” corresponds to the macrophage only.

**Figure S3:**
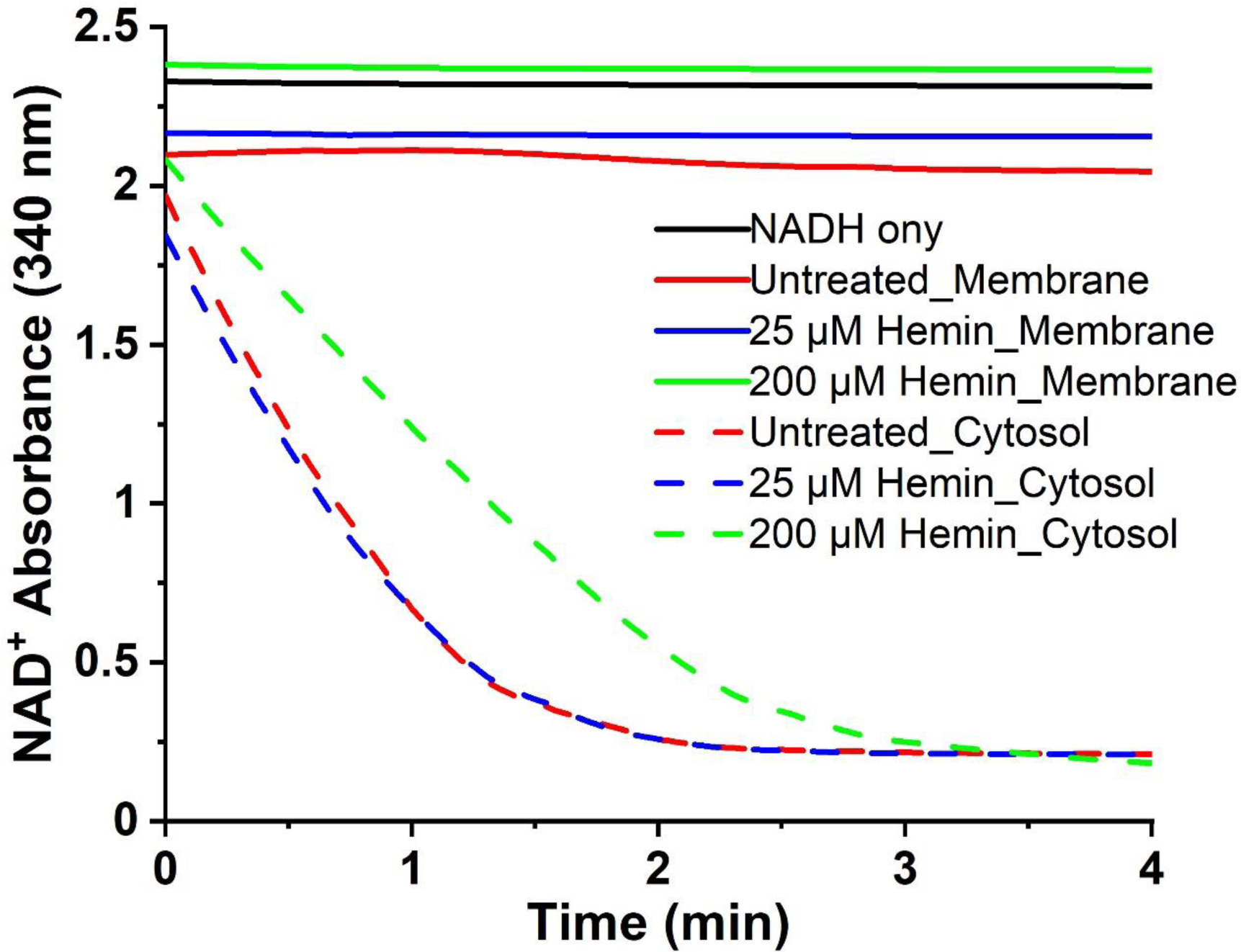
Purity assessment of membrane and cytosol fractions from macrophages. The macrophages either treated with hemin or remain untreated were homogenized using 2-bend 25 gauge needed and fractionated as described in methods section. The fractions are taken into a cuvette previously containing NADH + Sodium pyruvate mixture and read absorbance at 340 nm. The cytosolic fractions were showing reduction in NADH absorbance whereas the membrane fractions absorbance remain unchanged. The data shows the fractions were pure.

**Figure S4:**
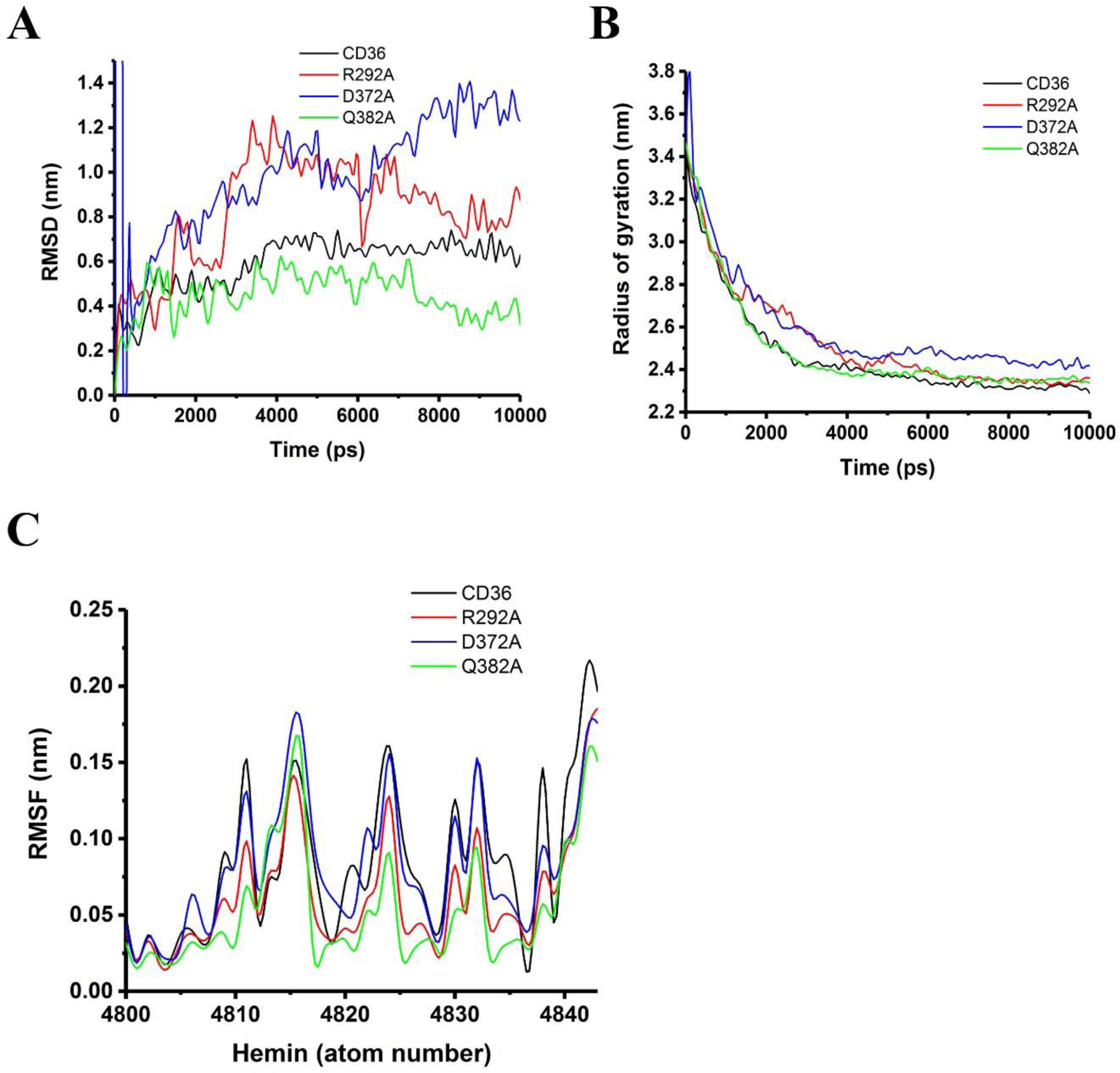
Stability analysis of wildtype and mutant proteins complexed with hemin using molecular dynamics simulations. The wildtype or mutant complexes were subjected to molecular dynamics simulations and the trajectories were analysed for the Root mean square deviation (RSMD), radius of gyration (Rg) and root mean square fluctuation (RMSF). **(A)** The root mean square deviation (RMSD) throughout the stimulation suggests the complexes are stable except the mutant D372A where a higher fluctuation were observed. Overall the complexes were converged stabilized during simulation. **(B)** The radius of gyration (Rg) of the CD36 and mutants indicates the compactness of the wildtype or mutants were unchanged upon ligand (hemin) binding. The simulation trajectory analysis suggests the proteins are tightly packed and remain in that condition throughout the simulation. **(C)** The RMSF of hemin atoms during simulation run was analysed using RMSF module of the Gromacs. There is no major perturbations were observed in hemin binding pocket as it evident from RMSF profile of hemin. The MD simulation analysis suggests the overall stability of wild type (CD36) and mutants complexed with hemin.

**Figure S5:**
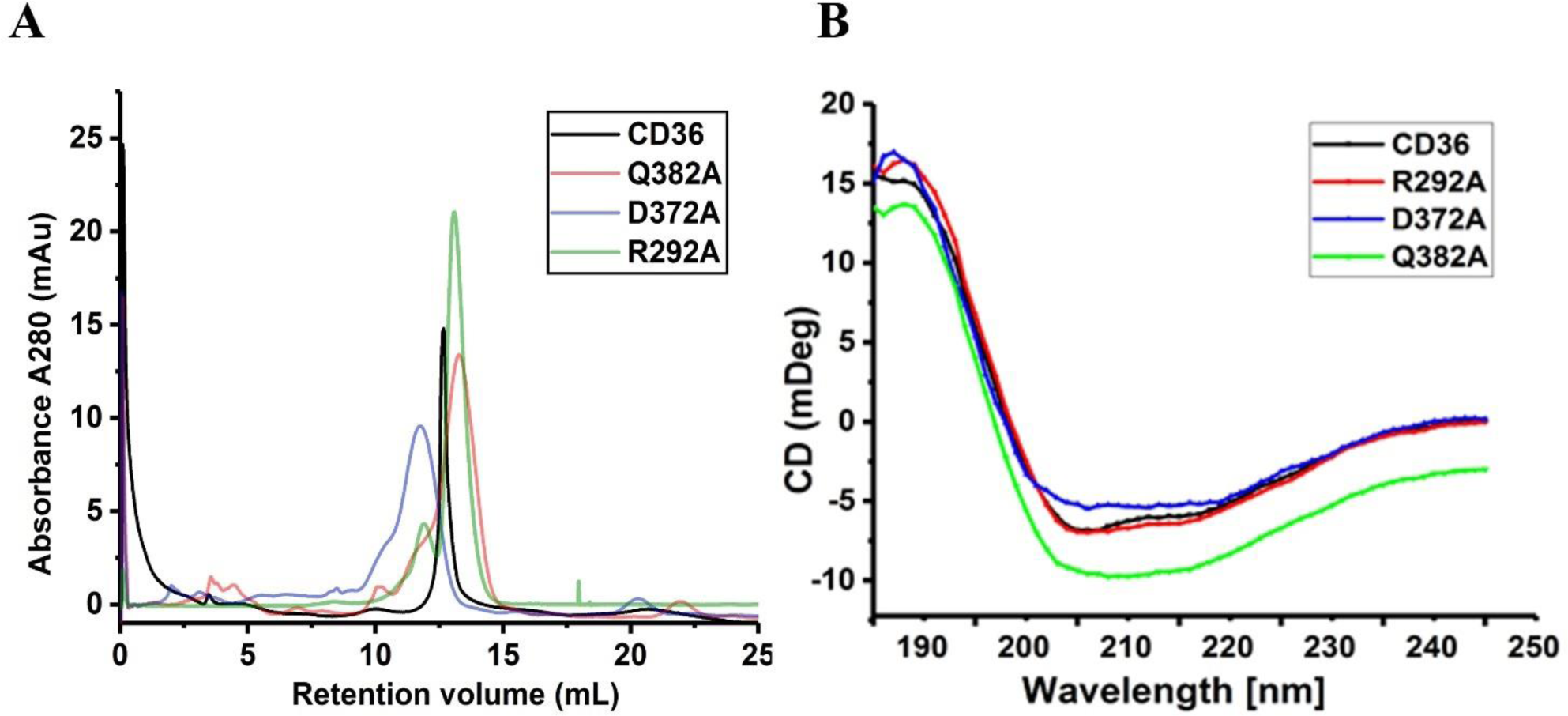
The mutants R292A, D372A and Q382A are monomeric in nature and structurally identical to the CD36. (A) The size exclusion chromatography profiling of the CD36 and mutants (R292A, D372A and Q382A) merged in same plot. The elution profile indicates there is no change in oligomeric status upon mutation to alanine. **(B)** Secondary structural characterization of CD36 and mutants using circular dichroism spectroscopy. The data indicates the mutants are structurally identical to the wildtype (CD36).

**Figure S6:**
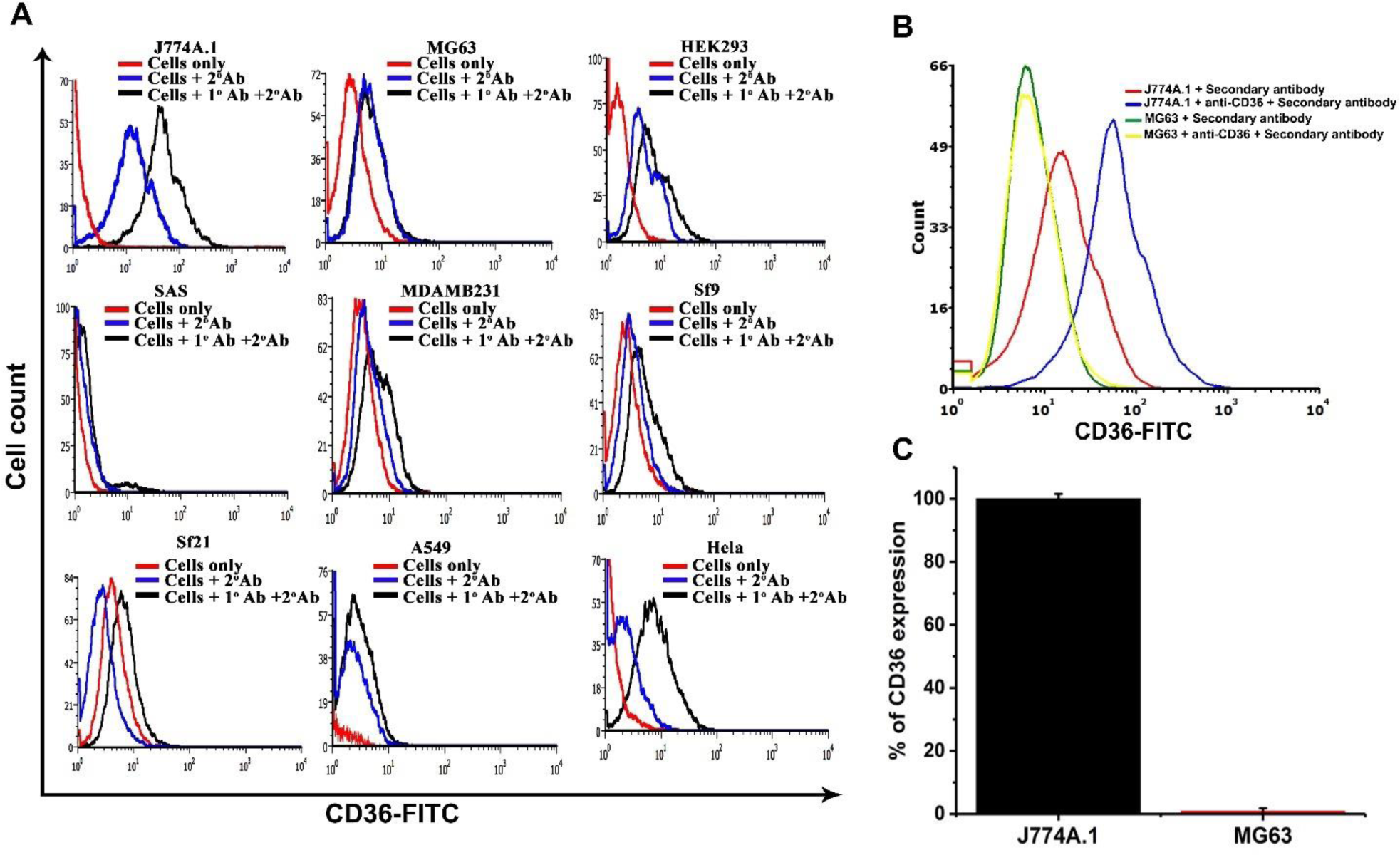
The CD36 expression in MG63 cells is minimal as compared to the J774A.1 cells. (A) The flow cytometry histogram representing the anti-CD36 antibody and FITC conjugated secondary antibody labelled J774A.1 and MG63 cells. The J774A.1 cells labelled with anti-CD36 antibody followed FITC conjugated secondary antibody showing fluorescence intensity mean of 60 whereas for MG63 it was 4. (B) The % of CD36 expression was semi quantified based on the flow cytometry data and plotted by considering the J774A.1 as 100%. The minimal expression was observed in MG63 cells. The data represents the result of three independent experiments.

**Figure S7:**
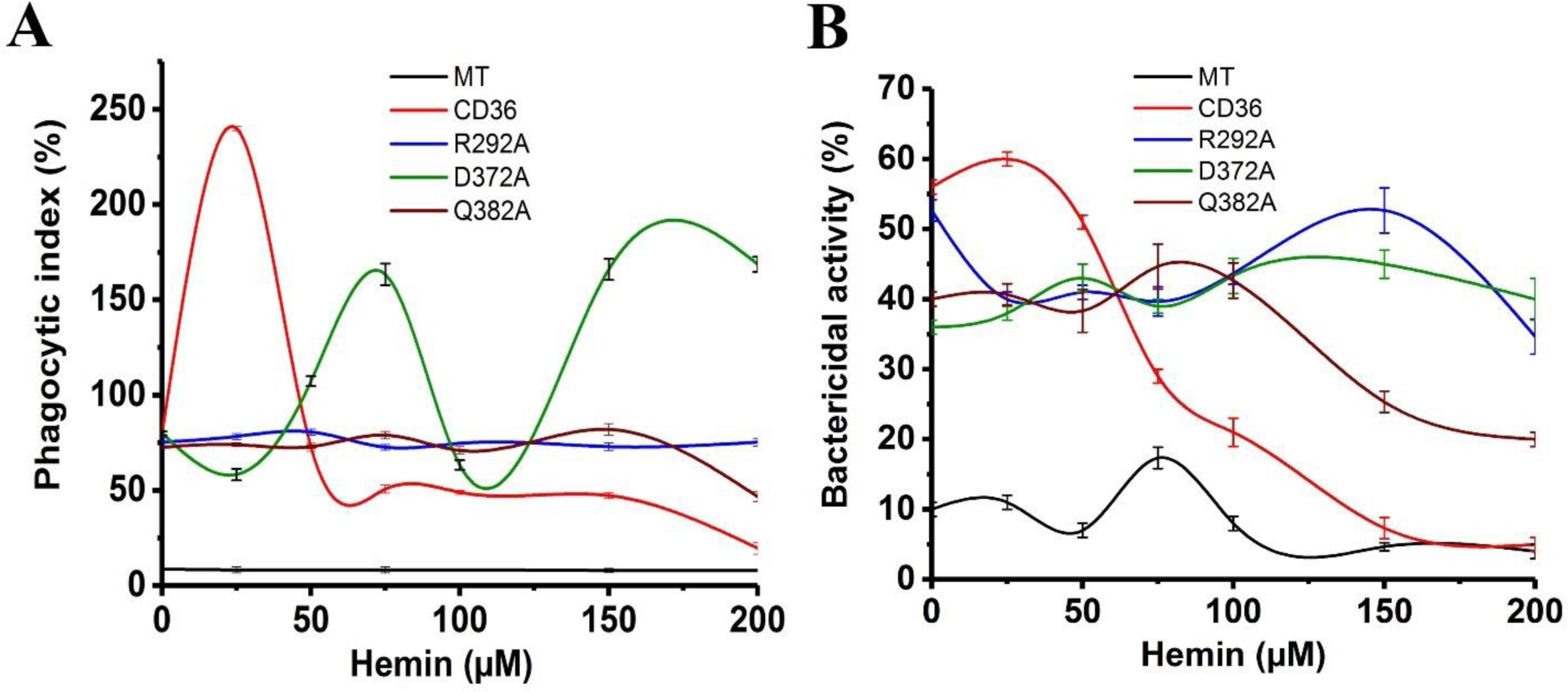
The residues R292, D372 and Q382 are crucial for hemin mediated immune-dysfunction. (A) The MG63 cells were transfected with either CD36 or its mutants. The successfully transfected cells either remain untreated or treated with 0-200 μM of hemin for 1 hr were incubated with FITC conjugated bacteria for 1 h. The cells were washed with PBS and the fluorescence counts were acquired in BD FACS calibur flow cytometer in FL-1 channel (λ_ex_-488 nm, λ_em_-530/30 nm). The phagocytic index was calculated as described in methods section. The CD36 transfected cells showing initial spike in phagocytosis index to 30% when compared to untreated and gradual decrease upon increasing concentration of hemin whereas R292A or Q382A transfected cells were showing no response to hemin stimulation. The D372A mutant showed abnormal phagocytosis activity upon hemin treatment. **(B)** The bactericidal activity of transfected MG63 cells assessed as described in methods section. The bactericidal activity of the CD36 transfected MG63 cells reducing with increasing hemin concentration but the decrease in activity is not prominent in mutants (R292A, D372A and Q382A) transfected MG63 cells.

**Figure S8:**
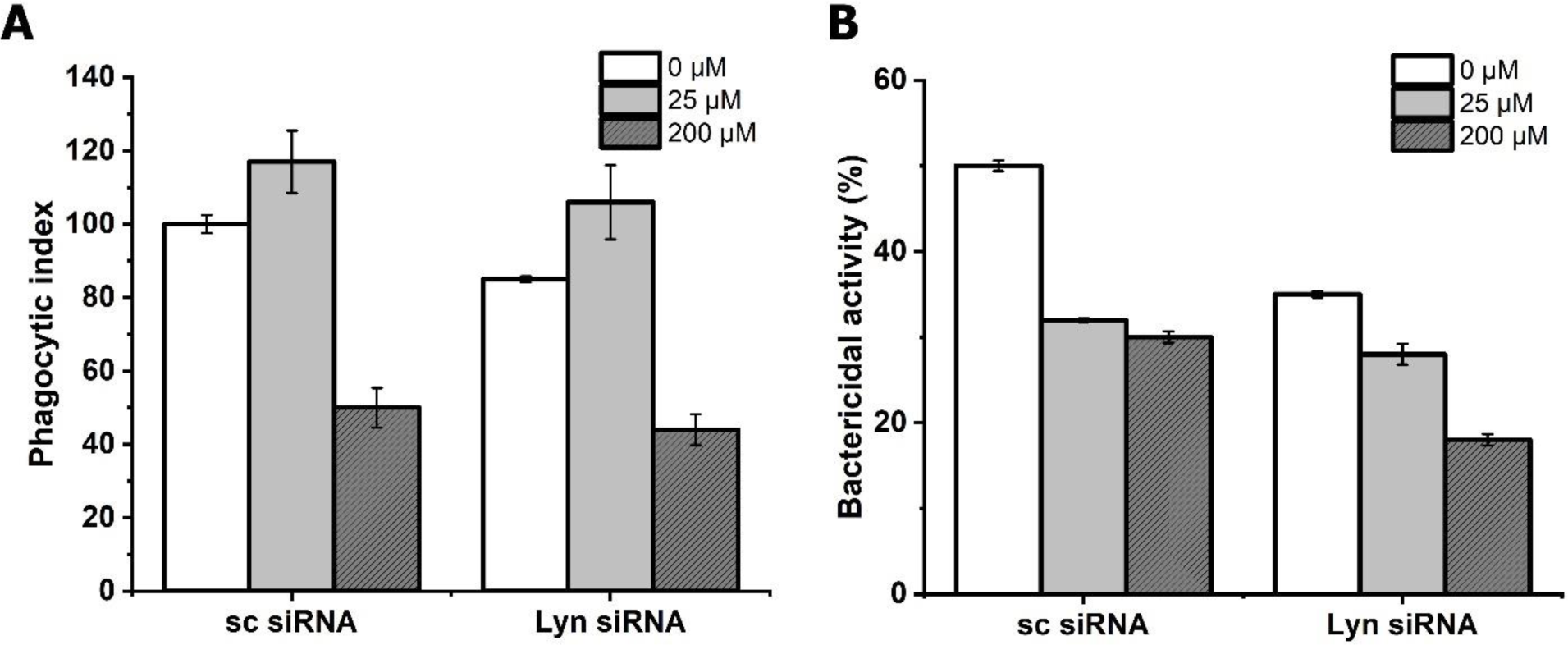
Assessment of phagocytic and bactericidal activity of Lyn knockdwon macrophages. The macrophages with Lyn siRNA transfected or control siRNA transfected and treated with hemin (25 or 200 μM) for 1 hr. Post treatment the phagocytic and bactericidal assays were performed as described in methods section. **(A)** The phagocytic activity of Lyn knockdown macrophages remained unchanged as compared to control siRNA. **(B)** Similar to phagocytosis assay, the bactericidal activity is also unaffected by lyn knockdown.

